# An experimental target-based platform in yeast for screening *Plasmodium vivax* deoxyhypusine synthase inhibitors

**DOI:** 10.1101/2023.11.23.568514

**Authors:** Suélen Fernandes Silva, Angélica Hollunder Klippel, Sunniva Sigurdardóttir, Sayyed Jalil Mahdizadeh, Ievgeniia Tiukova, Catarina Bourgard, Heloísa Monteiro do Amaral Prado, Renan Vinicius de Araujo, Elizabeth Bilsland, Ross D. King, Katlin Brauer Massirer, Leif A. Eriksson, Mário Henrique Bengtson, Per Sunnerhagen, Cleslei Fernando Zanelli

## Abstract

The enzyme deoxyhypusine synthase (DHS) catalyzes the first step in the post-translational modification of the eukaryotic translation factor 5A (eIF5A). This is the only protein known to contain the amino acid hypusine, which results from this modification. Both eIF5A and DHS are essential for cell viability in eukaryotes, and inhibiting DHS can be a promising strategy for the development of new therapeutic alternatives. The human and parasitic orthologous proteins are different enough to render selective targeting against infectious diseases; however, no DHS inhibitor selective for the parasite ortholog has previously been reported. Here, we established a yeast surrogate genetics platform to identify inhibitors of DHS from *Plasmodium vivax,* one of the major causative agents of malaria. We constructed genetically modified *Saccharomyces cerevisiae* strains expressing DHS genes from *Homo sapiens* (HsDHS) or *P. vivax* (PvDHS) in place of the endogenous DHS gene from *S. cerevisiae*. This new strain background was ∼60-fold more sensitive to an inhibitor of human DHS than the one previously used. Initially, a virtual screen using datasets from the ChEMBL-NTD database was performed. Candidate ligands were tested in growth assays using the newly generated yeast strains expressing heterologous DHS genes. Among these, two showed promise by preferentially reducing the growth of the PvDHS-expressing strain. Further, in a robotized assay, we screened 400 compounds from the Pathogen Box library using the same *S. cerevisiae* strains, and one compound preferentially reduced the growth of the PvDHS-expressing yeast strain. Western blot revealed that these compounds significantly reduced eIF5A hypusination in yeast. Our study demonstrates that this yeast-based platform is suitable for identifying and verifying candidate small molecule DHS inhibitors, selective for the parasite over the human ortholog.

## INTRODUCTION

Annually, approximately 250 million cases of malaria are reported, predominantly in low and middle-income countries, resulting in over 600,000 deaths [1]. Despite increased investments in malaria research and disease control over the past century, transmission rates remain high. Furthermore, the prognosis is expected to worsen due to the vector’s rapid range shift and its adaptability to urban environments, driven by climate change [2]. Some existing antimalarial drugs suffer from side effects and limited efficacy. Importantly, resistance to available antiplasmodial drugs is rapidly rising worldwide, emphasizing the urgent need for discovery of new and effective targets [3].

Efforts in malaria control have emphasized drug development targeting the responsible parasites, *Plasmodium spp*. The two main agents of malaria are *P. falciparum* and *P. vivax*, where *P. falciparum* accounts for the majority of malaria-related deaths. However, persistent *P. vivax* infections in the liver pose significant challenges to both treatment and disease elimination efforts, leading to high morbidity rates [4]. The shortage of *in vitro* culture methods for *Plasmodium* parasites presents challenges for antimalarial drug discovery. While continuous culture of *P. falciparum* in its intraerythrocytic stage is possible, there is currently no established continuous *in vitro* culturing system for *P. vivax*. This prompts the exploration of alternative approaches for assaying potential *P. vivax* drug target proteins, surrogate genetics systems in the yeast *Saccharomyces cerevisiae* emerging as an attractive option due to the facile genetic engineering in this organism [5–7].

Target-based drug discovery strategies have proven effective in developing new effective medicines against infectious diseases [8]. One promising target for investigation is the enzyme deoxyhypusine synthase (DHS). DHS catalyzes the first step of post-translational modification of the eukaryotic translation factor 5A (eIF5A). In this process, the aminobutyl group is transferred from the polyamine spermidine to a specific lysine residue in eIF5A (K50 in *H. sapiens* eIF5A) [9], forming deoxyhypusine. Subsequently, the enzyme deoxyhypusine hydroxylase (DOOH) hydroxylates deoxyhypusine, leading to the biosynthesis of the unique amino acid hypusine. Notably, eIF5A is the only protein identified to contain this distinctive amino acid [9,10].

Furthermore, both eIF5A and DHS are essential for cell viability in eukaryotes [11–13], including pathogens [14–16]. Hypusinated eIF5A plays a role in various pathologies and physiological processes, contributing to the regulation of cancer [17–20], immune-related diseases [21], diabetes [22,23], neurological disorders [24], aging [25], and the replication of certain viruses [26].

Small molecules capable of modulating mammalian DHS activity through distinct mechanisms, including competitive [27] and allosteric inhibition [28], have already been described. These known inhibitors are often considered as potential candidates for future anti-cancer therapies [29,30]. Given that DHS is an essential protein, it would be an attractive target for antiparasitic drugs, provided that such drugs would be sufficiently selective for the parasite ortholog of the protein. However, there is a notable absence of selective inhibitors described for DHS from parasites. This highlights the urgency to develop novel, targeted inhibitors specific for DHS from eukaryotic pathogenic organisms.

Target-based drug discovery campaigns typically rely on *in vitro* biochemical assays, which require protein purification and lack a cellular context [8,31,32]. Alternatively, designing systems with genetically modified *S. cerevisiae* strains can provide efficient platforms for target-based drug discovery within a eukaryotic cell [5–7,33–35].

In this work, we established a yeast target-based platform dedicated to the identification of novel inhibitors of *P. vivax* DHS. We genetically engineered isogenic *S. cerevisiae* strains to differ solely in their sole source of DHS, using either the orthologous gene from *P. vivax* or *H. sapiens*. Further enhancements were introduced into these strains to establish a robust system suitable for high-throughput screening, including genomic integration of genes encoding fluorescent proteins for precise quantification of cell proliferation. Additionally, genes related to pleiotropic drug resistance were deleted, to facilitate the entry of external molecules into the cells. Combined, these modifications resulted in a strain background ∼60-fold more sensitive to the human DHS inhibitor GC7. Using this system, we here identify three promising new small molecules targeting *P. vivax* DHS. These newly discovered candidate inhibitors for *P. vivax* DHS can serve as initial leads for the development of potent inhibitors against the *P. vivax* DHS protein. Moreover, the system we have established for *P. vivax* DHS will be applicable to DHS from other parasites and to other parasitic drug targets.

## RESULTS

### Homology modeling of *Plasmodium vivax* DHS

Currently, no 3D protein structure is available for *P. vivax* DHS (PvDHS). To further explore the structural features of *P. vivax* DHS and conduct *in silico* screening of small molecule libraries for potential enzyme inhibitors of this enzyme, we used the YASARA [36] software to generate homology models of PvDHS.

It is worth noting that all previously reported DHS structures are tetramers [7,37,38]. Moreover, DHS in *B. malayi* [7] and *H. sapiens* [28,37] are encoded by a single gene and exist as homotetramers. By contrast, DHS in trypanosomatids are heterotetramers, composed of two DHS paralogs [38]. Considering the structural similarities and the single-gene encoded PvDHS, we generated homotetramer models for this protein. Two models were selected for further investigation in our studies (Fig. S1).

The first model, PvDHS-model 1 (Fig. S1A), was constructed based on the deposited structure of *H. sapiens* DHS PDB ID: 6P4V [28], while the second model, PvDHS-model 2 (Fig. S1B), was based on *H. sapiens* DHS PDB ID: 6PGR [28]. We developed these two models (Fig. S1A and B) because 6P4V represents the structure in complex with the cofactor NAD^+^ and the competitive inhibitor GC7, while 6PGR is in complex with a ligand that induces a conformational change in which an α-helix is unfolded, forming a loop structure. Utilizing these models, our objective was to identify both competitive and/or allosteric inhibitors for PvDHS. SWISS-MODEL reported QMEAN scores for PvDHS-model 1 and PvDHS-model 2 were -2.62 and -3.45, respectively. In addition, the predicted local similarity based on the residue number revealed that residues from both binding sites were located in regions with estimated good quality (score ≥ 0.6) (Fig. S2).

The enzyme core of PvDHS shares many features with its human homologue. In fact, the PvDHS protein sequence exhibits a 59 % similarity with the human DHS (Fig. 1). Moreover, the residues in both the GC7 binding site and the allosteric site are conserved between HsDHS and PvDHS (as illustrated in Fig. 1). However, PvDHS and HsDHS differ in their overall length, PvDHS being 86 amino acids longer than HsDHS. Notably, these unique insertions are mainly located from the N-terminal to the middle of the protein extension (Fig. 1). The larger insertion (residues 224 to 263) aligns with residues that form an alpha helix in the human enzyme (α8 in Fig. 1). Currently, it remains unclear whether or how these extensions impact the PvDHS protein.

**Figure 1.**
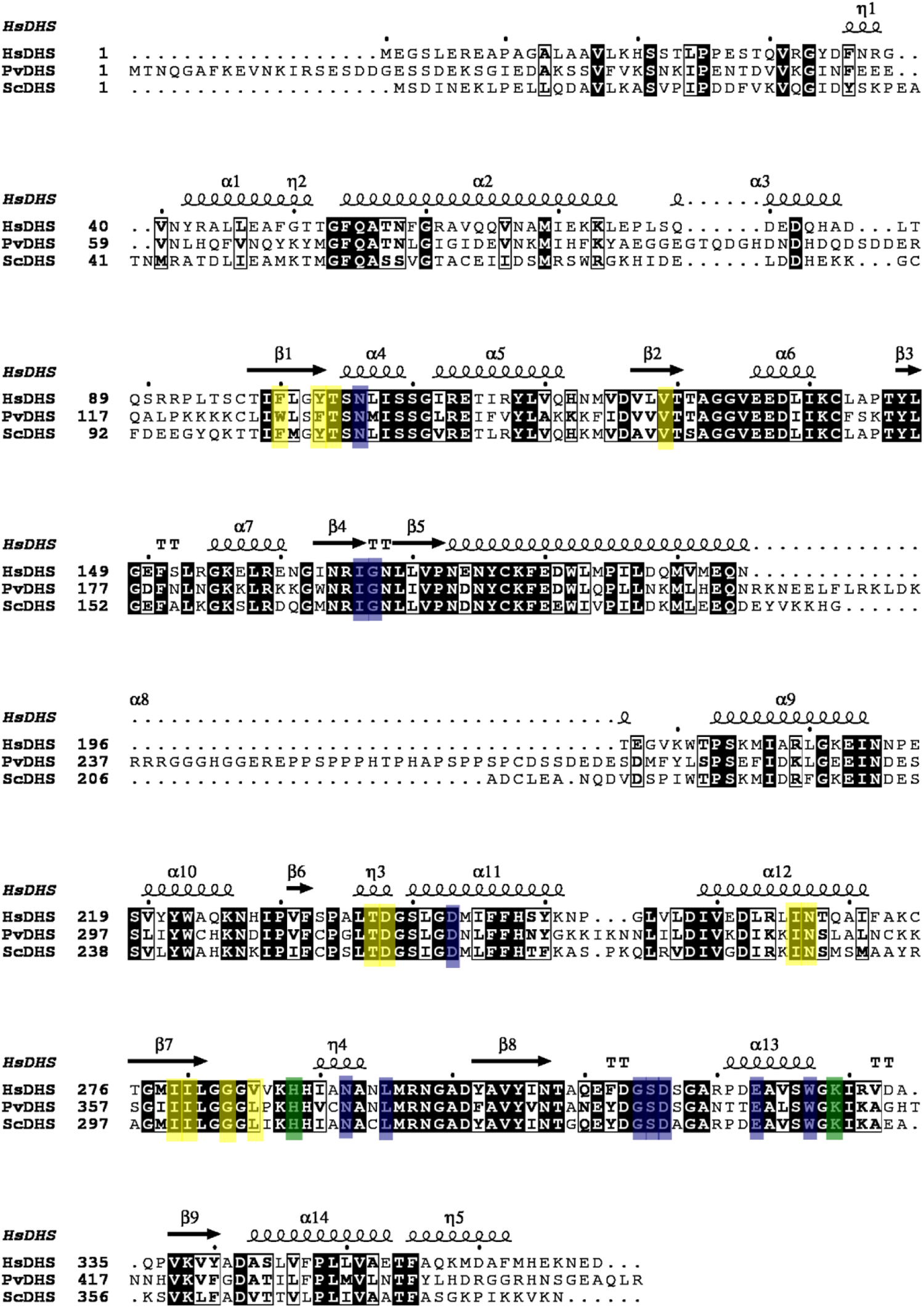
Structure-based sequence alignment of *H. sapiens* DHS (HsDHS), *P.vivax* DHS (PvDHS) and *S. cerevisiae* DHS (ScDHS). Residues indicated by a light-colored background participate in the orthosteric binding site (blue), allosteric binding site (yellow), or both (green). Background coloring for PvDHS and ScDHS follow that of the HsDHS proteins. Residues indicated by a black background or framed in a box are conserved in all DHSs analyzed here. The secondary structure (α-helices and β sheets) shown in the top line are for HsDHS. Protein sequences and structures used in the alignment were: HsDHS (UniProt ID P49366, PDB ID 6P4V), PvDHS (UniProt ID Q0KHM1) and ScDHS (UniProt ID P38791).

### Molecular docking

The compound library from the ChEMBL-NTD database (https://chembl.gitbook.io/chembl-ntd/) was chosen to conduct virtual screening of ligands toward PvDHS. This database contains several compound libraries that have been tested in previous campaigns for neglected tropical diseases (NTDs) and *Plasmodium* drug discovery campaigns. This library also contains information on the toxicity of some compounds in human cell culture. Moreover, there are studies reporting the identification of protein targets modulated by compounds from this repository [39,40].

Initially, the dataset was manually curated and filtered, primarily selecting sets that had been previously filtered for compounds tested and found to be active in phenotypic assays against parasites were chosen, except for dataset 23 which was not pre-filtered. The datasets chosen for this study are listed in the Zenodo repository. The molecular library prepared from the ChEMBL-NTD database comprised 212,736 structures, accounting for different possible protonation states for each compound.

Given the lack of specific inhibitors for *P. vivax* DHS, we predicted the binding sites based on co-crystallized ligands from human DHS structures (PDB IDs: 6P4V and 6PGR) [28], which served as the basis for the PvDHS models. We defined the binding residues for the known eukaryotic DHS spermidine mimic inhibitor, GC7 (PDB ID: 6P4V) [28], and of the human DHS allosteric ligand, 6-bromo-n-(1h-indol-4-yl)-1-benzothiophene-2-carboxamide (PDB ID: 6PGR) [28] to delineate the anchorage sites for the compounds to be screened. Molecular docking was performed in both the predicted binding sites of PvDHS, and the identified promising ligands were also docked into the HsDHS structures.

Before performing the virtual screening, we validated the robustness of the docking protocol by re-docking the co-crystallized ligands, GC7 and 8XY [28] into the human DHS structures (PDB IDs 6P4V and 6PGR, respectively) (Fig. S3). The docking poses of both compounds precisely aligned with the crystal structure, with RMSD values of 0.205 Å for GC7 and 0.208 Å for 8XY, respectively (Fig. S3). The corresponding docking scores were approximately –6.0 kcal/ mol for both compounds. As these ligands are known potent inhibitors of DHS (GC7 IC_50_ = 0.14 µM and 8XY IC_50_ = 0.062µM) [28,41], this value was used as a cutoff threshold to identify new promising ligands, as described in the Methods section.

Following the docking strategy (as described in the Methods section), we identified 100 and 85 promising hits (docking scores > – 6 kcal/mol) from the prepared structures in the ChEMBL-NTD database towards the orthosteric site (GC7 binding site) and allosteric site (8XY binding site) of PvDHS, respectively. The promising hits were also docked into the orthosteric or allosteric binding sites of the *H. sapiens* DHS protein structures, to assess the compound selectivity *in silico*.

Based on the docking score values in PvDHS compared to HsDHS, free energy of binding values, and commercial availability, 9 compounds (named N1 to N9), were selected as promising ligands for the PvDHS to be tested in yeast assays. The docking scores and free energy values for the ligands in both predicted binding sites of PvDHS and in HsDHS are listed in Table 1.

**Table 1.**
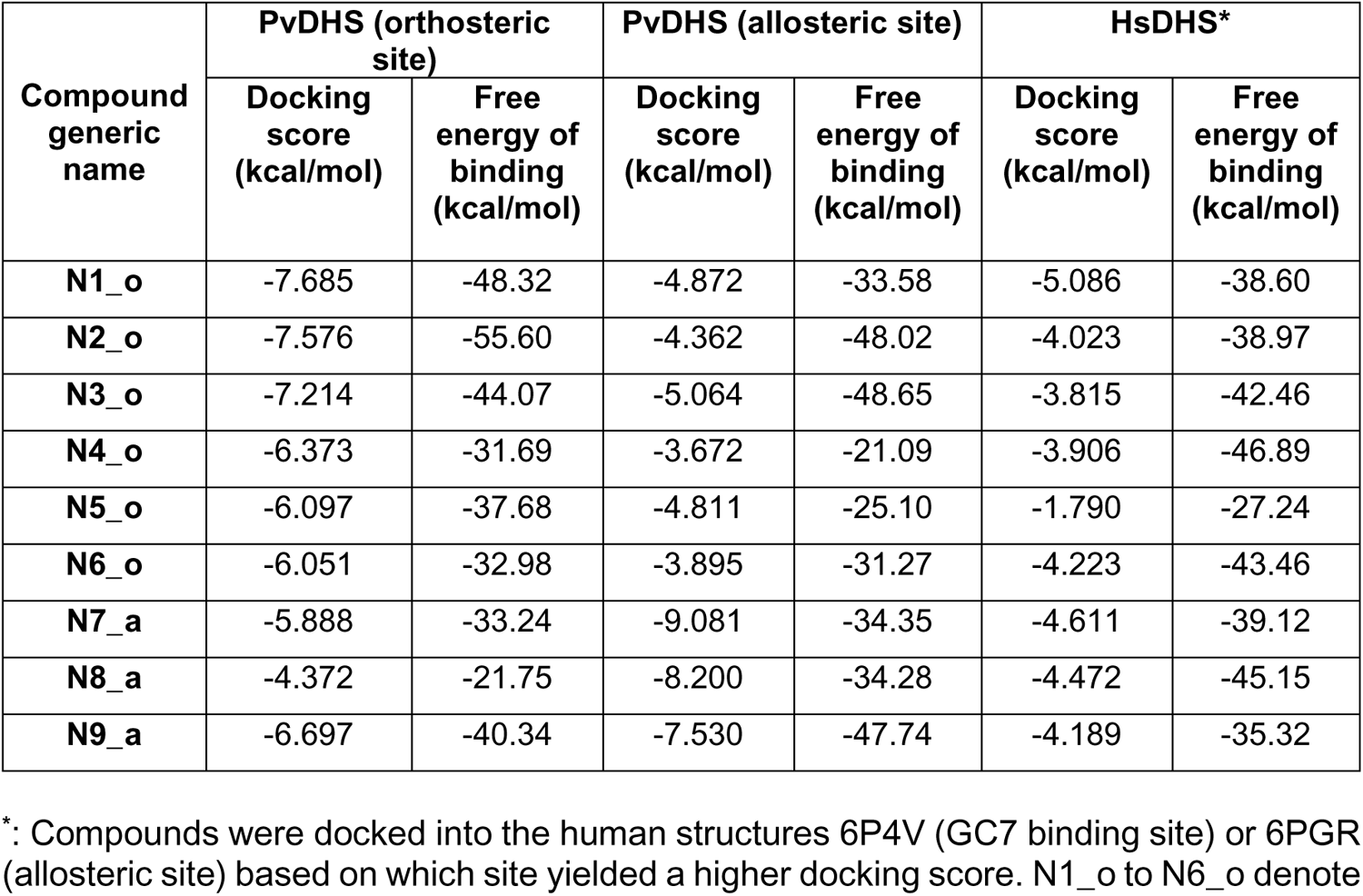
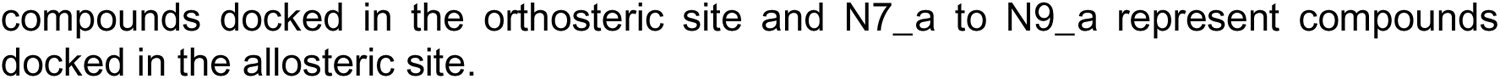
Docking score values obtained by molecular docking of the compounds in the PvDHS and HsDHS proteins.

| Compound generic name | PvDHS (orthosteric site) |  | PvDHS (allosteric site) |  | HsDHS* |  |
| --- | --- | --- | --- | --- | --- | --- |
|  | Docking score (kcal/mol) | Free energy of binding (kcal/mol) | Docking score (kcal/mol) | Free energy of binding (kcal/mol) | Docking score (kcal/mol) | Free energy of binding (kcal/mol) |
| N1_o | -7.685 | -48.32 | -4.872 | -33.58 | -5.086 | -38.60 |
| N2_o | -7.576 | -55.60 | -4.362 | -48.02 | -4.023 | -38.97 |
| N3_o | -7.214 | -44.07 | -5.064 | -48.65 | -3.815 | -42.46 |
| N4_o | -6.373 | -31.69 | -3.672 | -21.09 | -3.906 | -46.89 |
| N5_o | -6.097 | -37.68 | -4.811 | -25.10 | -1.790 | -27.24 |
| N6_o | -6.051 | -32.98 | -3.895 | -31.27 | -4.223 | -43.46 |
| N7_a | -5.888 | -33.24 | -9.081 | -34.35 | -4.611 | -39.12 |
| N8_a | -4.372 | -21.75 | -8.200 | -34.28 | -4.472 | -45.15 |
| N9_a | -6.697 | -40.34 | -7.530 | -47.74 | -4.189 | -35.32 |

\*: Compounds were docked into the human structures 6P4V (GC7 binding site) or 6PGR (allosteric site) based on which site yielded a higher docking score. N1\_o to N6\_o denote
compounds docked in the orthosteric site and N7\_a to N9\_a represent compounds docked in the allosteric site.

According to the docking results, compounds N1 to N6 (N1_o to N6_o in Table 1) were predicted to bind into the orthosteric site. As anticipated, similar to GC7 (Fig. S3), these compounds were positioned at the interface between subunits A and B of PvDHS (Fig. 2).

**Figure 2.**
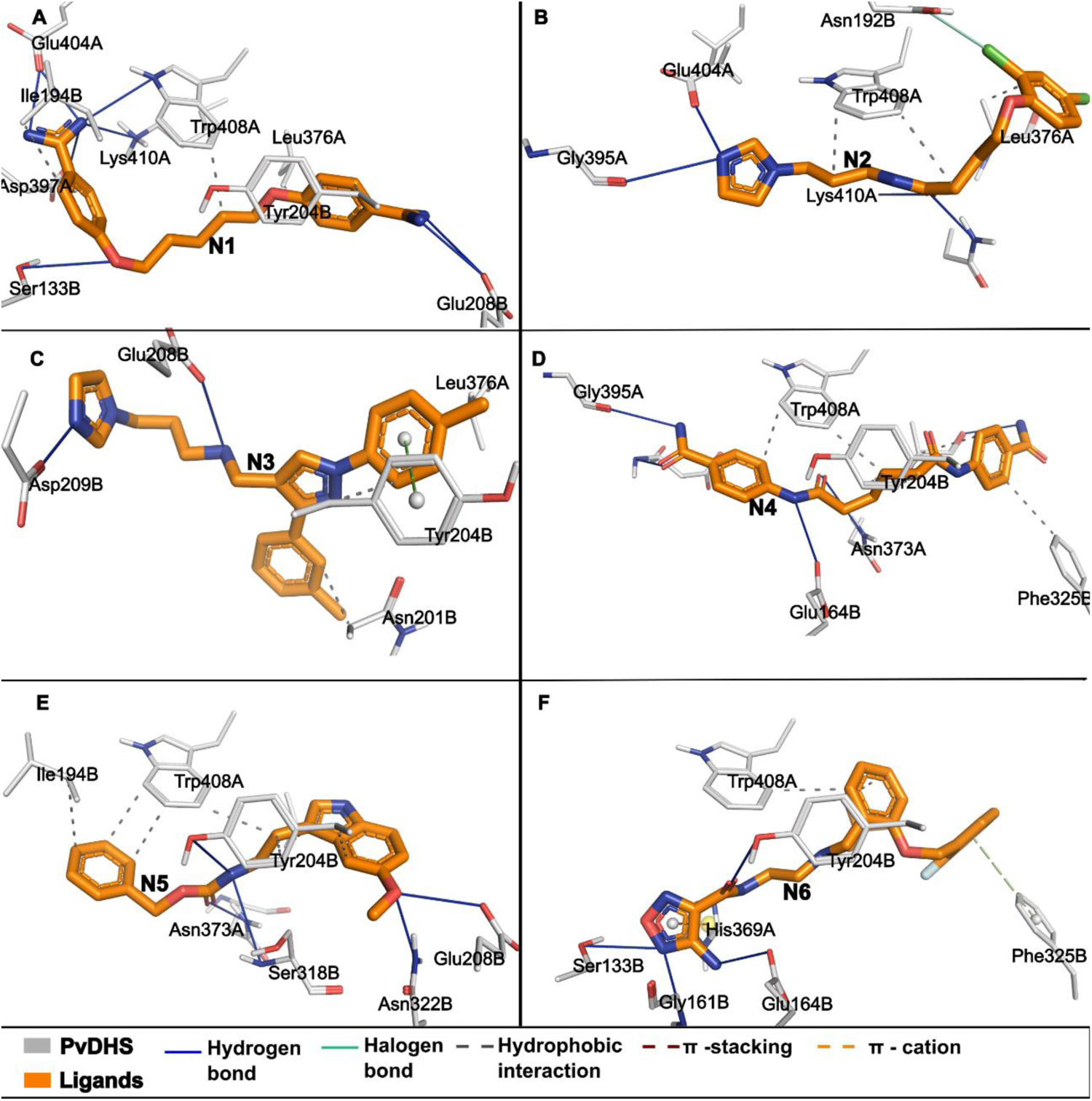
Predicted binding mode of the selected compounds in PvDHS. The binding mode and interaction profile of the selected compounds N1 to N6 (A to F, respectively; in orange, labeled in the figure) in PvDHS residues (gray). The residues from PvDHS that interact with the compounds are identified by the three-letter amino acid code followed by number and chain. Different type of interactions are represented by different colors and lines, and their corresponding meaning are described at the bottom of the figure.

The interaction profiles between the ligands N1_o to N6_o and the surrounding residues in subunits A and B within the active site of PvDHS are illustrated in Fig. 2. By contrast, compounds N7 to N9 (N7_a to N9_a in Table 1) were predicted to bind to the allosteric site of PvDHS (Table 1 and Fig. 3). The structures with the docked compounds are available in the Zenodo database.

**Figure 3.**
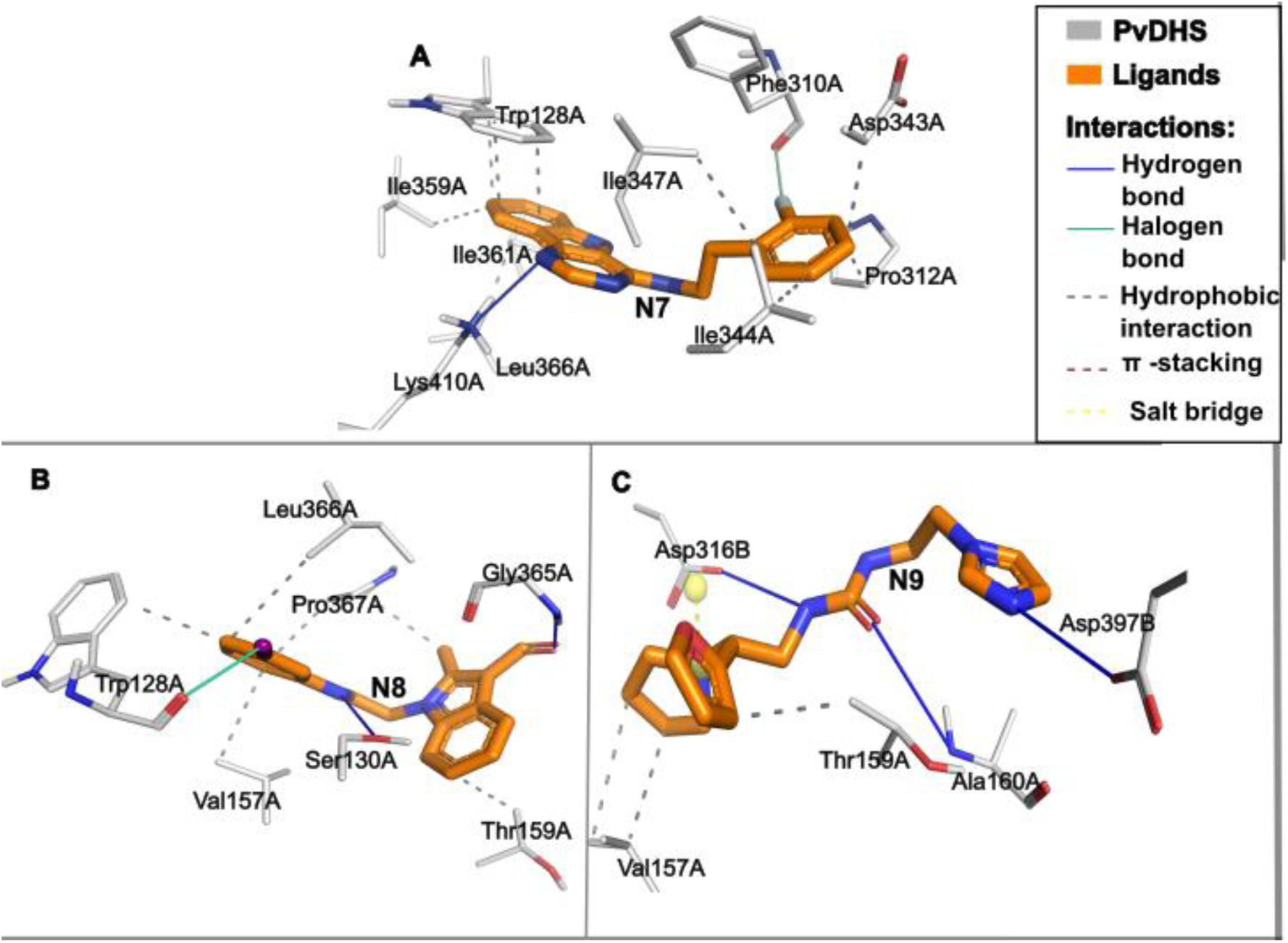
Predicted binding mode of compounds N7 to N9 in PvDHS. 3D views illustrating the interaction of compounds N7 to N9 (A to C, respectively; in orange, labeled in the figure) with PvDHS residues (gray). Residues from PvDHS that interact with the compounds are identified by three -letter amino acid code followed by number and chain. Different type of interactions are represented by various colors and dashes and their corresponding meaning described in the rectangle located in the right side of the figure.

### Establishment of an *S. cerevisiae* surrogate genetics platform for *P. vivax* DHS inhibitor discovery

The yeast *S. cerevisiae* shares conserved pathways with other branches of the eukaryotic phylogenetic tree, making it a valuable model for studying heterologous enzyme function [32,42]. A yeast surrogate genetics platform, involving the functional replacement of essential genes in yeast by heterologous genes, has proven effective for infectious diseases drug discovery [5,33,35]. Furthermore, parallel functional complementation with the human and parasite counterparts of DHS offers a way to pinpoint small molecules that selectively modulate the pathogen’s protein [33].

To establish a reliable yeast platform to search for *P. vivax* DHS inhibitors, we constructed a mutant *S. cerevisiae* strain with three primary genetic modifications: 1) deletion of a transporter (Snq2) and transcription factors (Pdr1, Pdr3) involved in pleiotropic drug response; 2) insertion of fluorescent markers (mCherry, Sapphire); and 3) substitution of the endogenous gene encoding ScDHS (*DYS1*) with orthologous genes expressing PvDHS or HsDHS from a chromosomal locus. Modifications 2 and 3 were performed using CRISPR-Cas9 techniques, as described in the Methods section. Integrating these modifications into the yeast platform strain’s genome is crucial, particularly for large-scale experiments. This ensures more consistent expression of the newly integrated gene, thereby enhancing the reliability and reproducibility of our assay strategy [43].

We performed the genetic modifications using a yeast strain with deletions of *PDR1*, *PDR3* and *SNQ2* (HA_SC_1352control [44], Table S2). *PDR1* and *PDR3* (<u>P</u>leiotropic <u>D</u>rug <u>R</u>esistance) encode redundant transcription factors positively regulating the expression of various membrane transporters involved in pleiotropic drug response, while *SNQ2* (Sensitivity to NitroQuinoline-oxide) encodes a plasma membrane ATP-binding cassette (ABC) transporter [45]. Deleting these genes simultaneously sensitizes yeast to a broad range of compounds, without compromising robustness [46].

For each strain, we inserted genes encoding different fluorescent proteins, mCherry or Sapphire, into the *CAN1 locus*, to serve as markers for cell abundance. This enables precise monitoring of growth through fluorescence and turbidity measurements. Additionally, chromosomal expression of the markers reduces noise and overcomes copy number variations, compared to expression from plasmids [47].

Subsequently, we replaced the endogenous essential gene encoding ScDHS (*DYS1*) with the orthologous DHS genes from *P. vivax* or *H. sapiens* in the genome. The deletion of ScDHS was confirmed by western blot, as no band was detected with the anti-Dys1 antibody for *S. cerevisiae* DHS (Fig. S4). ScDHS replacement was confirmed by DNA sequencing. Moreover, PvDHS and HsDHS expressed in yeast were capable of catalyzing eIF5A modification, hypusinating the endogenous yeast eIF5A (Fig. S4), which demonstrates the functional complementation of the strains.

### Validation of the yeast experimental platform for DHS inhibitor identification

Strains expressing DHS from *H. sapiens* or *P. vivax* (SFS04 and SFS05, respectively; Table S2) were tested for their ability to detect DHS inhibitors. To validate the platform, we conducted growth curves and western blot assays using GC7, a well-known inhibitor of human DHS [27].

It is important to note that in the newly generated strains, DHS expression was under transcriptional control of the *MET3* promoter. The *MET3* promoter can be repressed by adding methionine to the culture media [48], enabling tunable control of DHS expression. We determined the methionine concentration at which hypusine levels were similar in the PvDHS and HsDHS complemented strains, and used this information to define the assay conditions for testing the compounds selected by *in silico* screening.

The growth of the *dys1Δ* strain expressing HsDHS was analyzed in 0, 60 and 130 µM methionine, and the PvDHS strain growth was assessed in 0 and 7.5 µM methionine (Fig. S5). The strain expressing PvDHS did not grow in methionine concentrations exceeding 7.5 µM (data not shown). As expected from the greater evolutionary distance between *Plasmodium* and yeast, the level of hypusination in the strain complemented by PvDHS (approximately 40 % at 0 µM methionine, Fig. S5) was lower than in the strain with HsDHS (approximately 60 % at 0 µM methionine, Fig. S5), indicating that HsDHS complements the *dys1Δ* mutation more efficiently than does PvDHS. However, the hypusine levels in both strains were equal when HsDHS expression was regulated by 130 µM methionine (Fig. S5A). Thus, we established growth conditions of the pre-cultures at 130 µM methionine for HsDHS and 0 µM methionine for PvDHS.

It has previously been shown that reducing expression of the target protein in yeast sensitizes growth to inhibition by tested compounds [5]. Therefore, we performed a growth curve assay to examine the effect of GC7 on the human DHS-complemented strain under various methionine concentrations (ranging from 0 to 130 µM). Notably, we observed a significantly more pronounced growth reduction by GC7 at 130 µM methionine (Fig. 4A).

**Figure 4.**
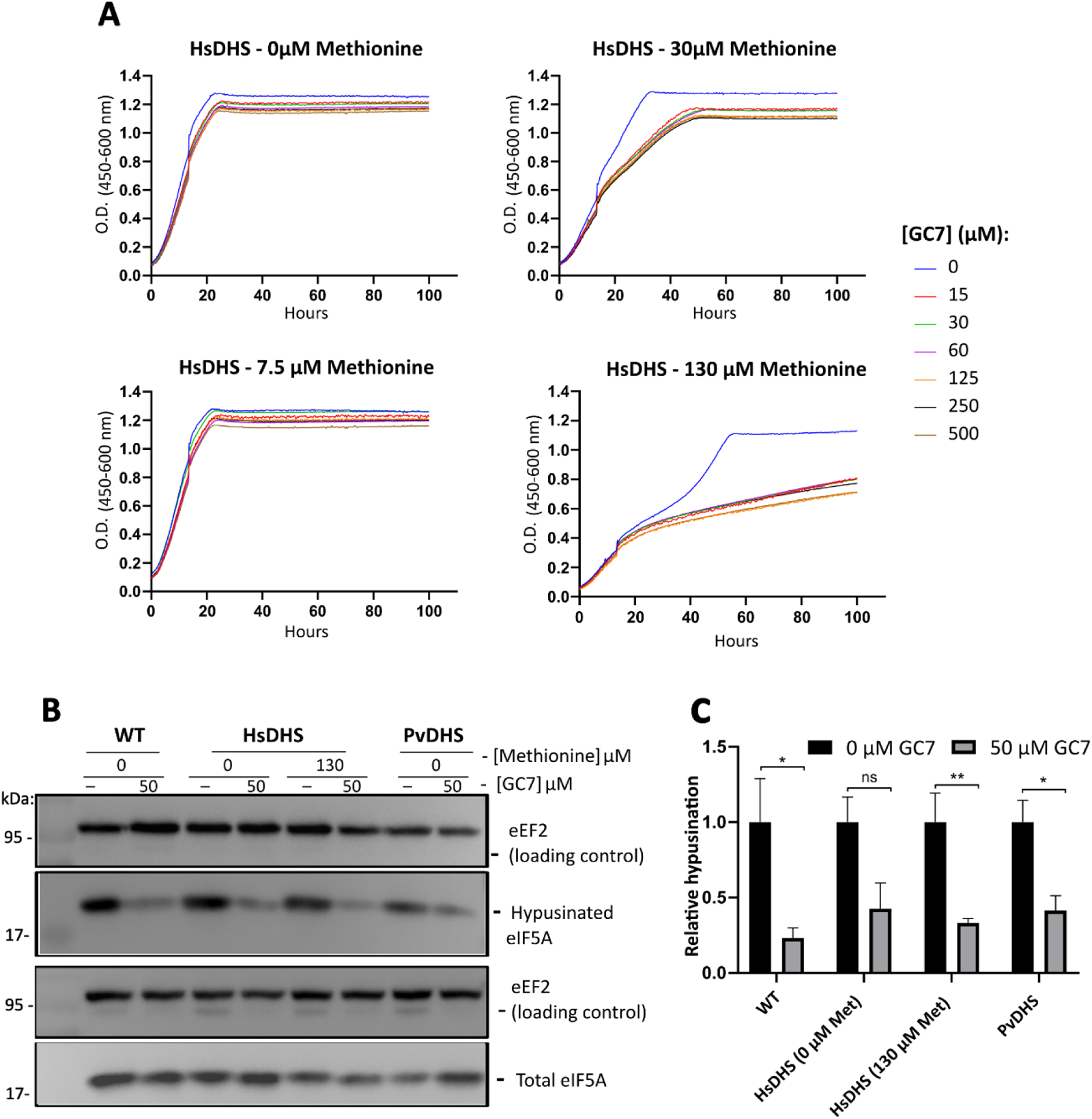
GC7 inhibits growth of *dys1Δ* yeast strain complemented by HsDHS. (A) Comparison of inhibition caused by different concentrations of GC7 (as indicated) in *dys1Δ* yeast complemented by HsDHS (SFS04, Table S2). The concentration of methionine used to regulate HsDHS expression is indicated in the figure. (B) Western blot detection of hypusination levels after 12 h of treatment with 50 µM GC7 in the wild-type and PvDHS and HsDHS-complemented strains. The methionine concentrations used in precultures (as mentioned in Material and Methods) are indicated in the figure. (C) Graph reporting the ratio between hypusinated eIF5A and total eIF5A under the conditions indicated in the figure. The bars represent mean of relative hypusination values ((hypusinated eIF5A/ eEF2) / total eIF5A/ eEF2)) ± standard deviation (n=3). A Student’s t-test was performed comparing the compound condition and the solvent control condition where an asterisk (*) indicates that the differences are statistically significant with 95 % confidence (p < 0.05); two asterisks (**) indicate that the differences are statistically significant with 99 % confidence (p < 0.01); three asterisks (***) indicate that the differences are significant statistically with 99.9 % confidence (p < 0.001), and ns indicates that the differences are not statistically significant (p > 0.05).

To evaluate if the observed phenotype was associated with a reduction in the final product of DHS, hypusine, we performed western blot analysis (Fig. 4B). In addition, the western blot was used to tested hypusine levels after addition of GC7 in the PvDHS-complemented strain, in the absence of methionine. This was defined based on the data presented in Fig. S5, which indicates that the hypusine levels are similar between HsDHS and PvDHS under these conditions. As shown in Fig. 4, treatment with 50 µM GC7 reduced hypusine levels (Fig. 4B and C).

### Yeast phenotypic assay using robotized platform

Using the Eve robotized platform programmed for incubation, agitation, and plate reading cycles [6], we tested the PvDHS inhibition of the nine compounds selected through *in silico* screening (see Table 1). Growth curve assays were performed (Figs. S6, S7 and S8) and metric parameters were extracted from each curve. The area under the curve was selected for comparison of different compound concentrations and the solvent control (DMSO) (Fig. 5).

In this *in vivo* assay, among the nine tested compounds two, N2 and N7, significantly reduced growth of the PvDHS-complemented strain compared to the strain complemented with HsDHS and the wild-type strain in the same genetic background (Figs. 5, S6, S7, and S8).

**Figure 5.**
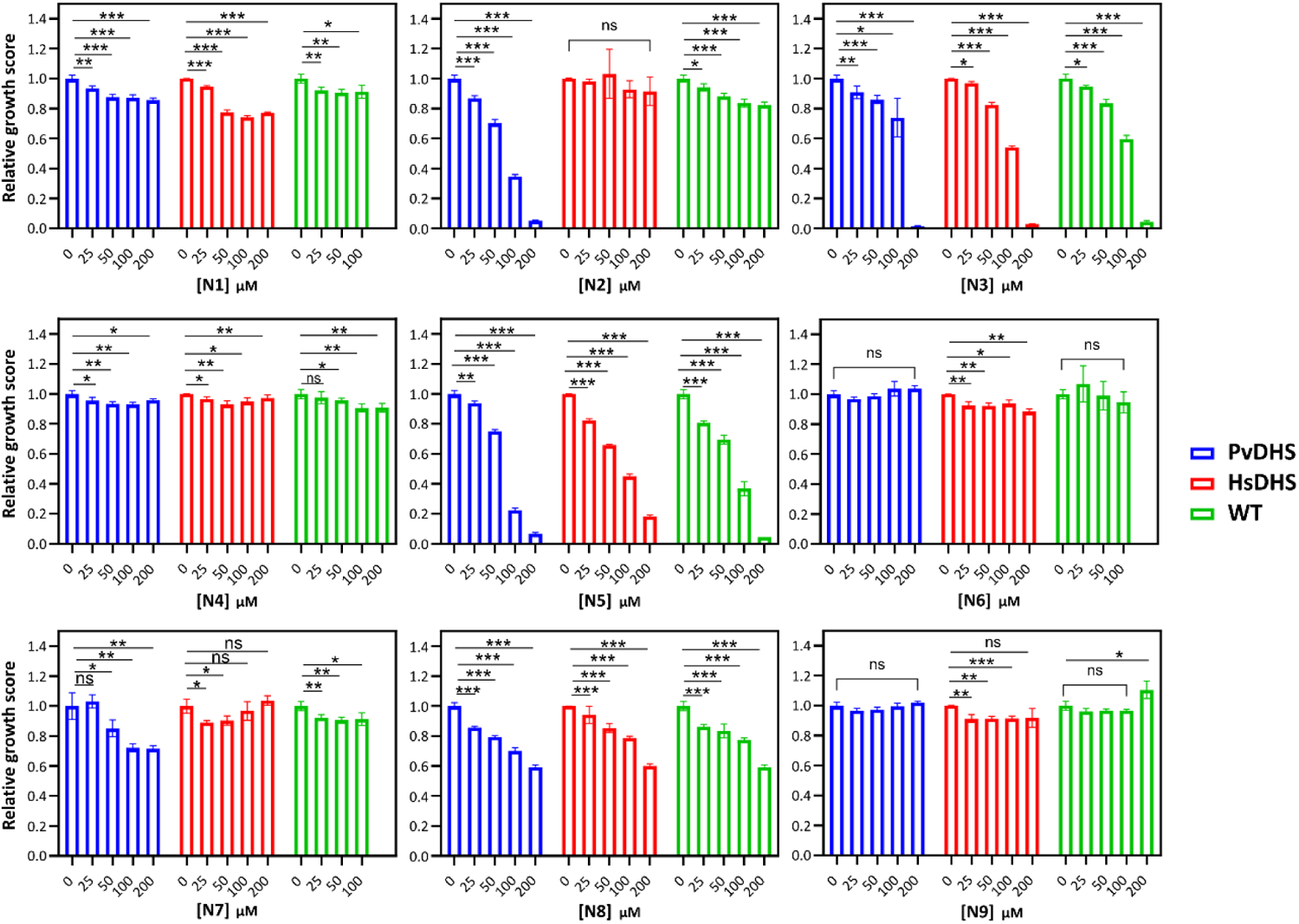
Detection of potential PvDHS selective inhibitors in the yeast-based assay. Bars represent the relative growth score of *S. cerevisiae dys1Δ* strains complemented by the indicated DHS enzymes (blue: PvDHS; red: HsDHS) and the relative growth score of the wild type (WT) isogenic strain is shown in green. The concentrations of the compounds range from 25 to 200 µM. Statistical significance levels indicated as in Fig. 4.

The 9 compounds were tested at concentrations ranging from 25 to 200 µM. When the growth curve was performed with the PvDHS-complemented strain, six compounds reduced the growth of this strain: N1, N2, N3, N5, N7 and N8. However, N1, N3, N5 and N8 reduced the growth of the *HsDHS* and WT strains at least to the same extent. In contrast, N2 and N7 reduced growth of the PvDHS-complemented strain more than the other strains (Figs. 5, S6, S7, and S8), indicating selectivity for PvDHS.

Notably, N2 exhibited dose-dependent inhibition of the PvDHS-expressing strain (Table S4), achieving a substantial 95 % reduction in growth at 200 µM, 65 % at 100 µM, 30 % at 50 µM, and 13 % at 25 µM of the compound (Fig. 5). In contrast, this compound did not significantly affect the growth of HsDHS (p > 0.05, represented by the red bars in Fig. 5). Furthermore, N2 at 200 µM only led to a modest 17 % reduction in the growth of the wild-type strain, expressing yeast DHS (illustrated by the green bars in Fig. 5). The selective growth reduction in the strain expressing PvDHS indicates that DHS is a major target of this compound.

Compound N7 (Table S4) resulted in a milder growth reduction phenotype, inhibiting approximately 30 % of the growth of PvDHS-expressing strain at 200 and 100 µM. In comparison, the strain expressing yeast DHS showed 9 % inhibition at 200 µM concentration, and no statistically significant inhibition was observed for the HsDHS-expressing strain (Fig. 5).

### Screening the Pathogen Box small-molecule library

In an alternative approach, using the same robotized phenotypic screen, we tested 400 compounds from the Pathogen Box small-molecule library (Medicines for Malaria Venture) at a concentration of 25 µM against the PvDHS-complemented strain (SFS05, Table S2). The majority of compounds did not affect the growth of the strains. However, 16 compounds (4 % of the total molecules in the library) reduced the growth of the PvDHS-complemented strain (Fig. S8). Not all of these 16 compounds were readily commercially available, so we proceeded with confirmatory tests using new batches of compounds PB1 and PB2 (Table S4). Out of these, we could only confirm growth inhibition of the PvDHS strain by PB1 in a follow-up growth curve assay; showing a dose-dependent response between 50 and 200 µM (Fig. 6). With the fresh compound, PB1 reduced 30% of the growth of the *dys1Δ::PvDHS* (Fig. 6) strain at 200 µM, and the inhibition being 12 % in the *dys1Δ* strain complemented with HsDHS (Fig. 6).

**Figure 6.**
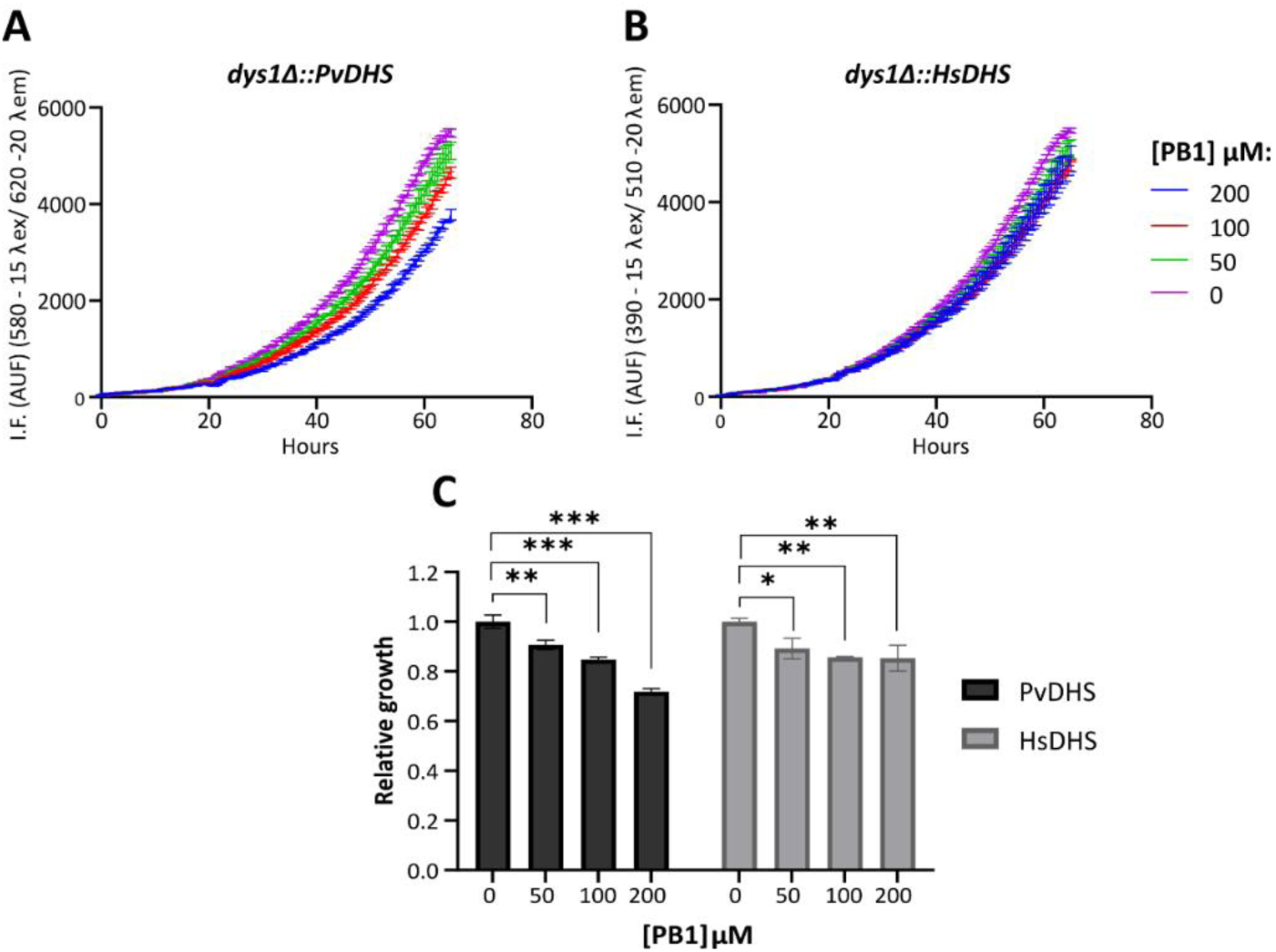
Relative growth of strains complemented by PvDHS and HsDHS in the presence of compound PB1. Growth curves were generated using different concentrations of PB1 (indicated in the figure) in the *dys1Δ::PvDHS* (A) and *dys1Δ::HsDHS* (B) strains. Each point represents the mean ± standard deviation of four experimental replicates. (C) Relative growth score, calculated by the curve with the compound in relation to the DMSO curve. Statistical significance levels indicated as in Fig. 4.

### Hypusination is reduced by N2, N7 and PB1 in cells expressing PvDHS, but not HsDHS

Finally, to verify whether the observed growth reduction phenotype in the yeast assay directly correlated with a decrease in the final product (hypusine) of the reaction catalyzed by DHS, we assessed the hypusination levels in the strains complemented with HsDHS, PvDHS, and the wild-type strain (SFS04, SFS05, and SFS01, respectively; Table S2) following a 12 h treatment with the compounds. Specifically, hits from the yeast screening, including N2 (at 100 µM), N7 (at 50 µM), and PB1 (at 100 µM) were tested for their ability to inhibit hypusination in eIF5A (Fig. 7).

Compounds N7, N2, and PB1 significantly reduced hypusination levels in the PvDHS-complemented strain (SFS05, Table S2) by 35 % (p < 0.05), 60 % (p < 0.05) and 70 % (p < 0.01), respectively (as shown in Fig. 7A and B). Importantly, no reduction in hypusination was observed in the other tested strains (WT and HsDHS) following a 12 h treatment with these compounds, indicating that the inhibition was specific for PvDHS.

**Figure 7.**
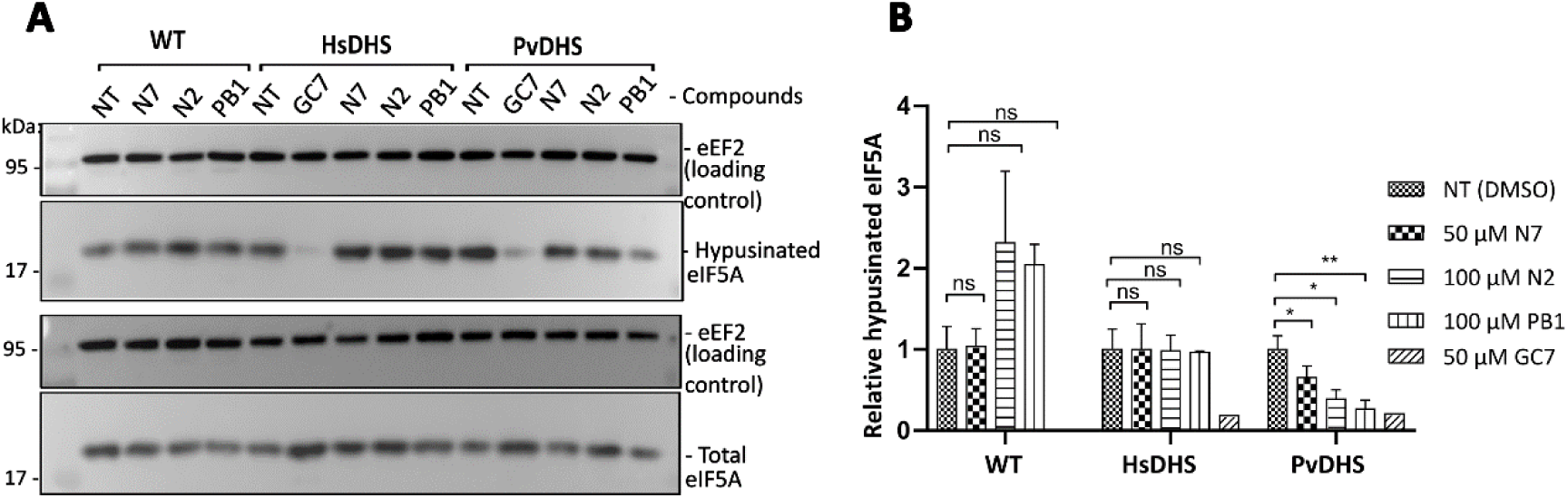
Hypusination of eIF5A in the presence of PvDHS inhibitor hits. (A) Detection by western blot of hypusination levels after 12 h of treatment with compounds indicated in the figure, at the following concentrations: GC7 (50 µM), N7 (50 µM) N2 (100 µM) and PB1 (100 µM). The methionine concentrations used in pre-cultures were 130 µM for HsDHS and 0 µM for PvDHS and WT. (B) Graph reporting the ratio between eIF5A hypusinated and total eIF5A under the conditions indicated in the figure. The bars represent the mean relative hypusination values ((hypusinated eIF5A / eEF2) / total eIF5A / eEF2)) ± standard deviation (n=3); except for GC7 where the bar represents the value for one experiment (n=1). A Student’s t-test was performed comparing the compound treated condition and the solvent control condition. Statistical significance levels indicated as in Fig. 4.

### Cytotoxicity

To assess the overall toxicity of N2, N7, and PB1, a metabolic viability assay was conducted in human cell cultures using the Presto Blue^TM^ cell viability reagent (ThermoFisher Scientific), relying on the redox reaction of resazurin. The compounds were tested on two types of human cells: breast cancer cells (MCF7) and hepatocarcinoma cells (HepG2).

Notably, compounds N2 and PB1 demonstrated no significant inhibition of cell viability at the tested concentrations (Fig. S10 A and B) indicating their potential for further drug development.

However, compound N7 exhibited a reduction in cell viability for the tested cell lines, with a predicted IC_50_ of 32 µM (see Fig. S10C). Previous studies, reported a 50 % growth inhibition (IC_50_) in Huh7 hepatocarcinoma cells at a concentration of 14 µM [49].

## DISCUSSION

We have here established a *S. cerevisiae* surrogate genetics system for high-throughput screening to identify new inhibitors of *P. vivax* DHS. This approach was an advancement of previous yeast surrogate systems designed for target-selective inhibitor searches, circumventing challenges associated with *in* vitro growth of parasites [5,34,35]. Utilizing CRISPR-Cas9 homology-directed repair, we have incorporated new features including fluorescent markers and genetic integration of the orthologous DHS genes, greatly enhancing the platform’s robustness and providing control over a wider range of target protein expression levels, using the weak regulatable *MET3* promoter.

Our new strain, with DHS expression regulated by the *MET3* promoter and reduced ABC exporter activity, exhibited improved sensitivity to compounds. This allowed growth inhibition at lower concentrations (15 µM) compared to previous studies using *Leishmania major* and *H. sapiens* DHS, where 1 mM GC7 was required due to poor compound permeability [7]. This corresponds to a 60-fold improvement of sensitivity.

Here, with this established yeast-based platform, we screened a total of 409 compounds against PvDHS, including 400 molecules from the Pathogen Box library (Medicines for Malaria Venture) and 9 compounds selected through virtual screening from a ChEMBL-NTD database. From this screening three compounds targeting PvDHS – N2, N7 and PB1 – were identified. A cytotoxicity assay in mammalian cell culture revealed minimal toxicity for N2 and PB1 in the tested cell lines.

Compounds N2 and N7 were previously tested against *P. falciparum* in an erythrocyte-based infection assay conducted by Novartis. N2 exhibited a potency of EC_50_ = 0.48 µM against *P. falciparum* 3D7 strain, and EC_50_ = 1.25 µM in the chloroquine-resistant W2 strain, whereas no cytotoxicity could be detected at 10 µM [49]. N7 exhibited a potency of EC_50_ = 0.15 µM in the *P. falciparum* W2 strain, and EC_50_ = 14 µM in the hepatocellular carcinoma cell line Huh7 [49], similar to our data from other human cell lines (Fig. S10). This indicates that despite its impact on cell viability, this compound was around 90 times more potent against *Plasmodium* cells, demonstrating its potential as an antimalarial agent.

Compound PB1 was previously recognized for its activity against *Mycobacterium tuberculosis* (MIC = 0.6 µM). PB1 is a dual-activity compound, *in vitro* targeting *M. tuberculosis* EthR (IC_50_ = 12 μM) and InhA (62 % inhibition at 100 μM) [50]. In fact, there is a co-crystal structure with this compound in complex with EthR, a transcriptional regulatory repressor protein (PDB ID: 5MXV) [50]. However, it is worth mentioning that the authors have raised concerns that this compound may have off-target effects or undergo pro-drug activation. This is suggested by the fact that the *in vitro* activity observed with these two proteins does not fully explain the low MIC exhibited by PB1.

We now propose DHS as a new protein target of PB1 in a different organism, *P. vivax*. We hope that future efforts in compound optimization can lead to improved target selectivity.

The distinctive amino acid hypusine is the final product of the reaction catalyzed by DHS in the post-translation modification of eIF5A [13]. Hence, we performed a secondary assay to investigate whether the growth reduction in yeast induced by these three compounds – N2, N7 and PB1 – was due to a decrease in the levels of hypusinated eIF5A. All three compounds significantly reduced the levels of *S. cerevisiae* hypusinated eIF5A levels selectively in the strain complemented by PvDHS (Fig. 7).

In conclusion, here we report the development of an alternative approach for drug discovery based on *S. cerevisiae* strains genetically engineered for increased sensitivity to external small molecules and more stable target gene expression, designed against the promising protein target, PvDHS. The identified compounds – N2, N7 and PB1 – show selective inhibition of PvDHS, demonstrating the potential of this yeast-based platform for antimalarial drug discovery.

## MATERIALS AND METHODS

### Plasmids and strains

Maintenance and cultivation of strains, preparation of culture media and solutions were conducted following standard protocols [51,52]. Plasmids and *S. cerevisiae* strains used in this study are listed in Tables S1 and S2, respectively. The sequence of primers used are given in Table S3.

### Synthetic compounds

The well-known DHS inhibitor, GC7 [27,29,53], and nine compounds (here named N1 to N9), selected from *in silico* screening in this work, were purchased from different suppliers (Table S4). Stocks of all compounds were prepared in DMSO at a concentration of 50 mM and stored at -20°C. The extended compound description as well as data about the potency against parasites from previous works are also listed in Table S4. Additionally, 400 small molecules from the Pathogen Box library (Medicines for Malaria Venture) were tested. The stocks of these compounds were at a concentration of 10 mM in DMSO (Sigma) and stored at -80° C. New batches of two compounds (named PB1 and PB2, Table S4) from this collection were acquired for confirmatory testing.

### Virtual screening of the ChEMBL-NTD repository

The 3D structures of the DHS protein from *P. vivax* were generated through homology-modelling using the YASARA software [36] and protein sequence retrieved from UniProt database (full length sequence; UniProt ID Q0KHM). Protein models quality were estimated using SWISS-MODEL Structure assessment online platform [54]. For *H. sapiens* DHS, high-resolution crystallographic structures were readily available (PDB IDs: 6P4V and 6PGR [28]), which were utilized in the screening campaigns. The protein structures underwent preparation with the Protein Preparation Wizard (Schrödinger Release 2021-2: Schrödinger Suite 2021-2 Protein Preparation Wizard) [55] within the Maestro software (Schrödinger Release 2021-2: Maestro, NY, 2021), applying default parameters. Protein preparation included removing water molecules and ligands, addition of hydrogens, assigning bonds and bond orders, completing missing loops and/or side chains using Prime [56,57], and optimizing hydrogen bonding network by adjusting the protonation states of Asp, Glu and tautomeric states of His to match a pH of 7.0 ± 2.0. Subsequently, geometric refinements were conducted using the OPLS4 force field [58] in restrained structural minimization.

Ligand structures were acquired from the ChEMBL-NTD database (https://chembl.gitbook.io/chembl-ntd/) in SDF format, and prepared using the LigPrep tool (Schrödinger Release 2021-2: Schrödinger Suite 2021-2, LigPrep) [55]. Ionization and protonation states were assigned in the pH range of 7.0 ± 2.0 using Epik [59] and energy minimization was performed with the OPLS4 force field [58].

Coordinates corresponding to the amino acids involved in the binding to the DHS inhibitor GC7 [13] and the allosteric inhibitor [28] were defined to generate the docking grid-boxes in the prepared protein structures. Molecular docking was executed using the Virtual Screening Workflow (Schrödinger Release 2021-2: Schrödinger Suite 2021-2, Virtual Screening Workflow Glide) [60], in the following selection stages: top 10 % poses selected by Glide HTVS (high-throughput virtual screening) were used in the Glide SP (standard precision); top 10 % poses from Glide SP were used in Glide XP (extra precision); and post-processing employing Molecular Mechanics with Generalized Born and Surface Area (MM-GBSA) method for the free energy of binding calculation using Prime [61].

In this strategy, we initially re-docked the co-crystalized ligands into the corresponding receptor protein upon which the PvDHS models were based. The docking score values of the co-crystalized ligands were employed as a cutoff threshold to consider new compounds as promising hits. Finally, we analyzed the free energy of binding values and the spatial disposition of the ligands within the binding sites to predict whether the compound would fit well within the receptor site.

### Construction of an *S. cerevisiae* DHS surrogate genetics platform

We developed a yeast surrogate genetics platform by adapting previously described CRISPR/Cas9 methodologies [47,62]. *S. cerevisiae* strains with two primary modifications were created: chromosomal integration of sequences expressing the fluorescent proteins mCherry and Sapphire, and replacement of the essential gene *DYS1* (which encodes DHS in *S. cerevisiae*) with DHS genes from *P. vivax* or *H. sapiens*. These genes were integrated into the *DYS1* locus in the genome of the *S. cerevisiae* working strain.

The genetic background for generating these strains was the *S. cerevisiae* HA_SC_1352control strain [44] (Table S2). This strain has deletions of *PDR1* and *PDR3*, encoding transcription factors that regulate pleiotropic drug responses, as well as *SNQ2*, a membrane transporter of the ABC family [46].

### Genomic integration of mCherry and Sapphire genes

The coding sequences for the fluorescent proteins Sapphire and mCherry, under control of the *TDH3* constitutive promoter, were inserted in the *S. cerevisiae* HA_SC_1352control strain into the *CAN1* locus, using CRISPR/Cas9-mediated homology-directed repair [62].

### sgRNA design, assembly and preparation

The design, assembly, and preparation of the single guide RNA (sgRNA) targeting *CAN1* locus were adapted from the Benchling platform (https://benchling.com/pub/ellis-crisprtools). We selected a previously reported sgRNA targeting *CAN1* [62] (Table S3). The oligonucleotides containing the guide sequences were annealed and cloned into the *BsmB*I site of the sgRNA entry vector pWS082 (Table S1) using Golden Gate assembly protocol [63]. This process replaced the GFP coding sequence from pWS082, generating the pWS082-CAN1 sgRNA construct (Table S1). Half of the reaction mixture was used for transformation into *E. coli* 10-beta^TM^ (New England Biolabs). White colonies (indicating the absence of GFP) were selected, and the pWS082-CAN1 sgRNA plasmid (Table S1) was extracted using the GeneJet Plasmid miniprep kit (ThermoFisher Scientific). Accuracy of the cloning was confirmed through DNA sequencing.

### Yeast transformation

We used a standard yeast transformation protocol [64] incorporating three DNA components: 300 ng of pWS082-CAN1 sgRNA vector, linearized by *EcoR*V digestion; 100 ng of the pWS172 previously digested with *BsmB*I and 3 µg of PCR-amplified donor DNA. The donor DNA consisted of the mCherry/Sapphire sequences, with the addition of *CAN1* homologous arms. This process generated the yeast strains SFS01 and SFS02 (Table S2).

### Deletion of DYS1 and replacement by DHS from H. sapiens and P. vivax

The coding sequences of DHS from *H. sapiens* (Gene ID: 1725, isoform A, here referred to as HsDHS) and *P. vivax* (Gene ID: 5476136, here referred to as PvDHS) were synthesized with codon usage optimization for expression in *S. cerevisiae* (Text S1) and amplified by PCR using the oligonucleotides described in Table S3. Subsequently, they were cloned into the *BamH*I-*Pst*I sites of the yeast expression vector pCM188-MET3 (Table S1), resulting in the generation of the following constructs: pCM188-MET3-HsDHS and pCM188-MET3-PvDHS (Table S1). For details on all constructs generated in this study, including primer sequences, consult Tables S1 and S3. The DNA constructs were used in this work for genomic integration protocols and for the yeast-based functional complementation system, and were verified by sequencing.

### sgRNA design, assembly and preparation

The sgRNA targeting *DYS1* was designed using the online platform ATUM (https://www.atum.bio/eCommerce/cas9) (Table S3). The assembly and preparation process followed the same protocol described for *CAN1* in this work, resulting in construction of the plasmid pWS082-DYS1 sgRNA.

### Yeast transformation

The transformation reaction followed the same procedure as the integration of the fluorescent proteins into the *CAN1* locus. However, the DNA components included in the reaction were as follows: 300 ng of pWS082-DYS1 sgRNA vector, linearized by *EcoR*V digestion; 100 ng of the pWS158 previously digested with *BsmB*I; and at least 5 µg of PCR-amplified donor DNA. The donor DNA comprised the MET3pr-DHS sequences from *H. sapiens* or *P. vivax-CYCt* cassette, with the addition of *DYS1* homologous arms. These linear fragments were transformed into strains SFS01 and SFS02 (Table S2), generating strains SFS05 and SFS04 (Table S2), respectively. To confirm genome integrations, colony PCR was conducted using a forward primer annealing upstream of the *DYS1* locus and the other primer annealing the CDS of the newly inserted gene (Table S3), as previously described [65], and DNA sequencing.

### Culture conditions

The *S. cerevisiae* strains used in this work are described in Table 1. Synthetic defined (SD) medium (0.19 % yeast nitrogen base without amino acids and ammonium sulfate, 0.5 % ammonium sulfate, 2 % glucose) with dropout of the appropriate amino acids was used for culture maintenance and assays performance.

### Validation of the platform in the presence of the drug GC7

Growth curves of yeast strains expressing DHS from *H. sapiens* (Strain SFS04, Table S2) or *P. vivax* (strain SFS05, Table S2) were analyzed in the presence and absence of inhibitors. Prior to testing the inhibition by new compounds, we conducted an initial assay standardization using the well-known inhibitor of the human DHS enzyme, GC7 [27]. The assay was done in the presence of GC7 and different methionine concentrations added to the SD medium. With that, we aimed to evaluate GC7 inhibition across various levels of DHS expression.

To prepare for the experiment, a 5 mL pre-culture was grown at 30°C under agitation until it reached approximately 1x10^7^ cells/mL. Once this density was achieved, the pre-cultures were diluted to a concentration of approximately 1x10^6^ cells/mL (OD_600nm_ = 0.2). The cultures were then incubated under agitation for approximately 3 h until they entered the exponential growth phase (OD _600nm_ ∼0.6). Just before the experiment, the cultures were diluted to OD_600nm_ = 0.1 in the appropriate SC medium and dispensed in 200 µL aliquots per well in 100-well plates (Honeycomb plates, Labsystems Oy) using the automated OT2 robot pipettor (Opentrons). The strains were then cultivated for 72 h at low agitation, with absorbance values (OD_450–600nm_) measured every 20 min using the Bioscreen C plate reader equipment (Oy Growth Curves Ab Ltd).

### Phenotypic assays using a robotized platform

The strains were cultivated in SD liquid medium, utilizing the optimal methionine concentrations determined through the growth validation assay.

Cultures with a final OD_600nm_ of 0.1 were dispensed into black 384-well plates (Greiner) using the Multidrop Combi reagent dispenser system (ThermoFisher). The compounds selected for virtual screening (Table S4) were transferred to the plates using the Bravo Liquid Handling platform (Agilent) to final concentrations ranging from 25 to 200 µM for each compound (resulting in a final DMSO concentration of 1.25 %) in a total volume of 80 µL.

The cultures were incubated at 30°C, and their growth was monitored as previously described [6]. An automated platform was programmed for cycles of incubation, agitation, and plate reading using the Eve robot [6]. Fluorescence values (mCherry = 580 nm λ_excitation_ / 612 nm λ_emission_; Sapphire = 400 nm λ_excitation_ / 511 nm λ_emission_) and absorbance (O.D._600nm_) were measured every 20 min over a period of 3 days using the BMG Polarstar plate reader. The experiments were conducted in two different experiments, each in quadruplicate.

To visually assess growth reduction, growth curves were plotted using the mean of fluorescence values or absorbance, along with their respective standard deviations, derived from quadruplicate measurements. A solvent control sample was included in the same graph as a reference.

A model was fitted, considering logistic growth model equations. The parameters relative to the curves were extracted using the Growthcurver package [66] in the R studio software (Version 2022 12.0 for Windows, Posit Software). The area under the curve (AUC) values were extracted for each curve and selected to calculate a relative score (relative growth = AUC compound / AUC DMSO). Student’s t-tests were conducted for four-sample comparisons, comparing the growth in the presence of compound with growth in solvent-only control.

In a different approach, compounds from the Pathogen Box library (Medicines for Malaria Venture) were also tested in the strain complemented by PvDHS with the same methodology. The experiment was carried out in quadruplicate, in one experiment. The curves representing the means were plotted in the R software. Two candidates from this screen were selected for confirmation (Table S4).

### Evaluation of hypusination levels by western blot assay

The hypusination levels of the complemented strains were evaluated by western blot. The strains were cultured for approximately 14 h in SD medium without methionine. Pre-cultures were then diluted to OD_600nm_ = 0.15 and grown during 10 h in fresh SD medium with different methionine concentrations. Cells were harvested by centrifugation at 4°C and stored at -80°C.

For the assays in the presence of inhibitors, after 10 h of cell growth in SD medium with the specified methionine concentration, cells were washed and diluted again to OD_600nm_ = 0.15 in SD medium lacking methionine. Inhibitor candidates were added at a concentration of 100 µM, except for compounds GC7 (inhibition control) and GNf-Pf-3738, which were added at a concentration of 50 µM. A control for 0 % inhibition was also prepared, containing only cells and the DMSO vehicle. After 12 h of treatment, cells were collected by centrifugation at 4°C and stored at -80°C.

Cell pellets were resuspended in 40 μL of lysis buffer (200 mM Tris/HCl, pH 7.0; 2 mM DTT; 2 mM EDTA; 2 mM PMSF) and lysed by agitation for 10 min with glass beads. The cells were clarified by centrifugation at 20,000 x g for 15 min at 4°C, and the supernatant was collected. Protein quantification was performed using the BCA protein quantification assay (QPro BCA kit, Cyanagen) against a bovine serum albumin (BSA) calibration curve (Sigma Aldrich).

A total of 12 µg of total protein was loaded onto a 12 % SDS-PAGE gel and transferred to a methanol pre-activated PVDF membrane using a semi-dry transfer system (Bio Rad, catalog number 1704150) following the equipment’s standard protocol. The western blot employed the following antibodies: rabbit polyclonal anti-eIF5A (yeast) antibody at a 1:10,000 dilution; rabbit polyclonal anti-hypusine (Millipore) antibody at a 1:2,000 dilution; and a rabbit polyclonal anti-eEF2 (yeast) antibody at a 1:25,000 dilution (sample loading control) (ThermoFisher), followed by a secondary anti-rabbit antibody at a 1:20,000 dilution (Sigma Aldrich).

The proteins of interest were detected using an enhanced chemiluminescence detection system (ECL, Amersham) on the Uvitec photodocumentation system (Alliance). The bands detected in the Western blot were quantified using GelAnalyzer 19.1 software [67]. The bands detected for anti-eIF5A and anti-hypusine were normalized in relation to the loading control bands (anti-eEF2), and the hypusination rate was calculated by dividing the values obtained for hypusinated eIF5A by the values quantified for total eIF5A.

### Evaluation of cytotoxicity in mammalian cell lines

#### Cell culture conditions

The tumorigenic cell lines MCF7 (breast adenocarcinoma, ATTC: HTB-22) and HepG2 (hepatocellular carcinoma, ATCC: HB-8065) were cultivated in DMEM (Gibco^TM^ DMEM, ThermoFisher Scientific) culture media supplemented with 10 % fetal bovine serum (Gibco^TM^ FBS, ThermoFisher Scientific) and 1 % penicillin-streptomycin (PenStrep, ThermoFisher Scientific). The cell lines were incubated at 37°C with 5 % CO_2_.

#### Evaluation of cytotoxicity using resazurin assay

Cell viability in the presence of compounds was assessed using the resazurin-based cell viability reagent, Presto Blue^TM^ (Thermo Fisher Scientific). MCF7 and HepG2 cells were seeded into 96-well plates (TPP) at a concentration of 10^4^ cells/well in 200 µL of culture medium and incubated for approximately 24 h until the cells adhered to the plate. The compounds selected as DHS inhibitor candidates in the yeast assay were added to the plates at concentrations ranging from 0.1 to 100 µM (with a maximum DMSO concentration of 0.4 %) and incubated for 24 h. After this, 10 % Presto Blue^TM^ was added, and the cells were further incubated for 1 h. Resofurin fluorescence was measured at an excitation wavelength of 544 ± 15 nm and an emission wavelength of 590 ± 15 nm using the spectrofluorometer POLARstar Omega (BMG Labtech). Each experiment was conducted in duplicate, and the experiments were repeated twice.

Cell viability percentages were determined by the quotient between the fluorescence intensity values of cells treated with the compounds by the values from solvent control cells, multiplied by 100. The concentrations that inhibited 50 % of cell growth (IC_50_) were calculated through non-linear regression. The percentage of cell viability was plotted against the logarithmic concentration of the compounds using GraphPad PRISM software (version 8.0 for the Microsoft Windows operating system), and the results are represented as the mean ± standard deviation.

## ACKNOWLEDGEMENTS

We thank the Medicines for Malaria Venture for the gift of a Pathogen Box, and Hanna Alalam for the yeast strain HA_SC_1352control. The Swedish national infrastructure for computing (SNIC) is gratefully acknowledged for allocation of computing time on the cluster Vera at C3SE and Tetralith at NSC. This work was supported by grants from the Swedish Research Council (2016-05627, 2019-03684, and 2021-03667), and the Swedish Foundation for International Cooperation in Research and Higher Education (BR2018-8017). SFS was the recipient of a fellowship from CAPES PRINT-Brazil (88887.570728/2020-00) and from CNPq (141241/2019-5 and 380882/2023-0). AHK was the recipient of PhD fellowships from CAPES (8887.161266/2017-00), CNPq (380893/2023-1), and from FAPESP (2018/16672-1 and 2019/24812-0). HM and RV were the recipients of fellowships from FAPESP (2021/04867-5 and 2022/10512-8, respectively). KBM and MHB were recipients of an INCT grant (CNPQ 465651/ 2014-3 /FAPESP 2014/50897-0).

## Supplementary Figures

**Figure S1.**
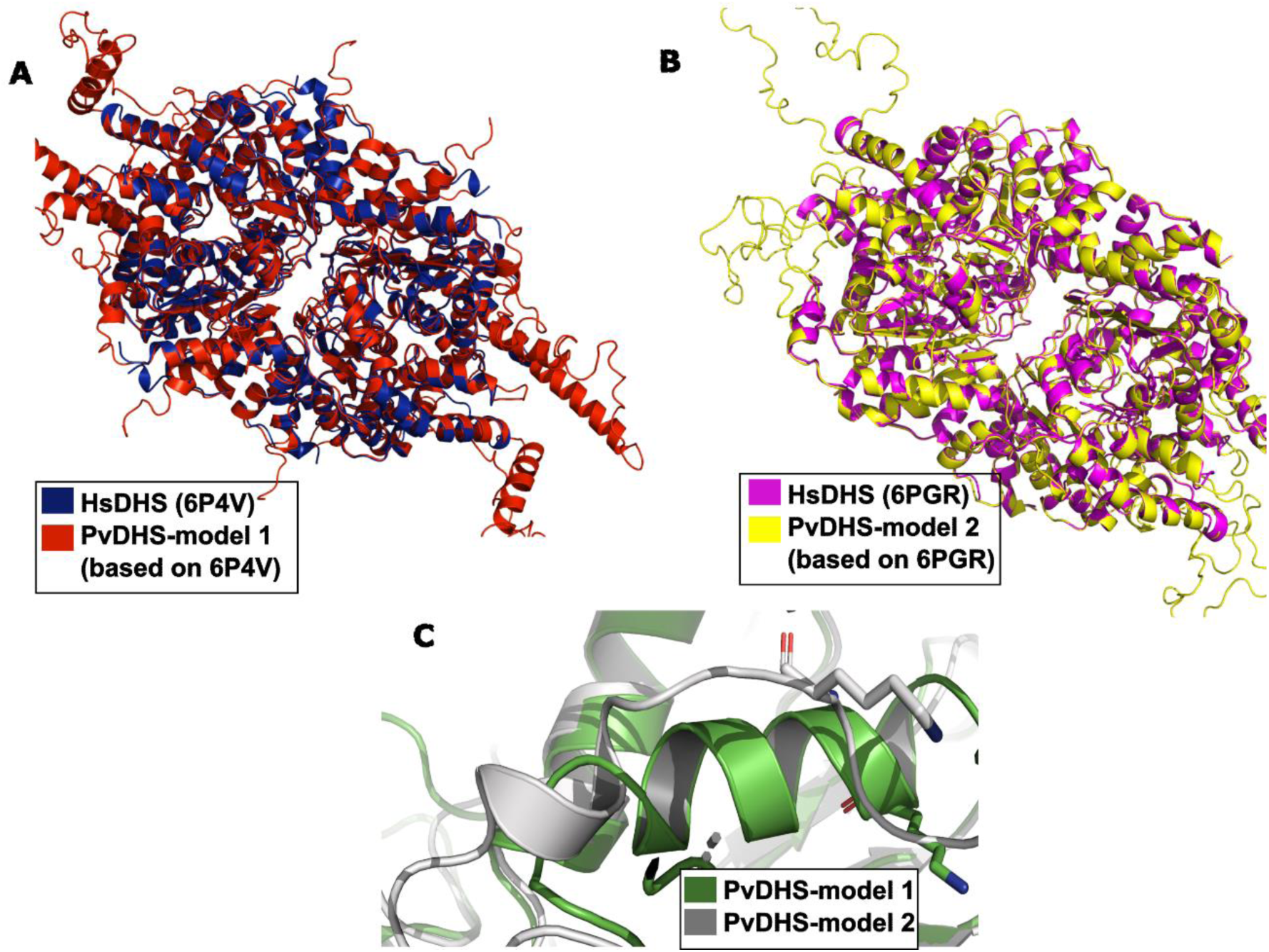
Homology models of *P. vivax* DHS homotetramer. (A) Cartoon representation of the PvDHS – model 1 (red) superposed on the HsDHS structure 6P4V [28] (blue). (B) Overlay of PvDHS – model 2 (yellow) superposed on the HsDHS structure 6PGR [28] (magenta). (C) Overlay of α-helix unfolded in PvDHS-model 2 (gray) with the corresponding position in PvDHS-model1 (green).

**Figure S2.**
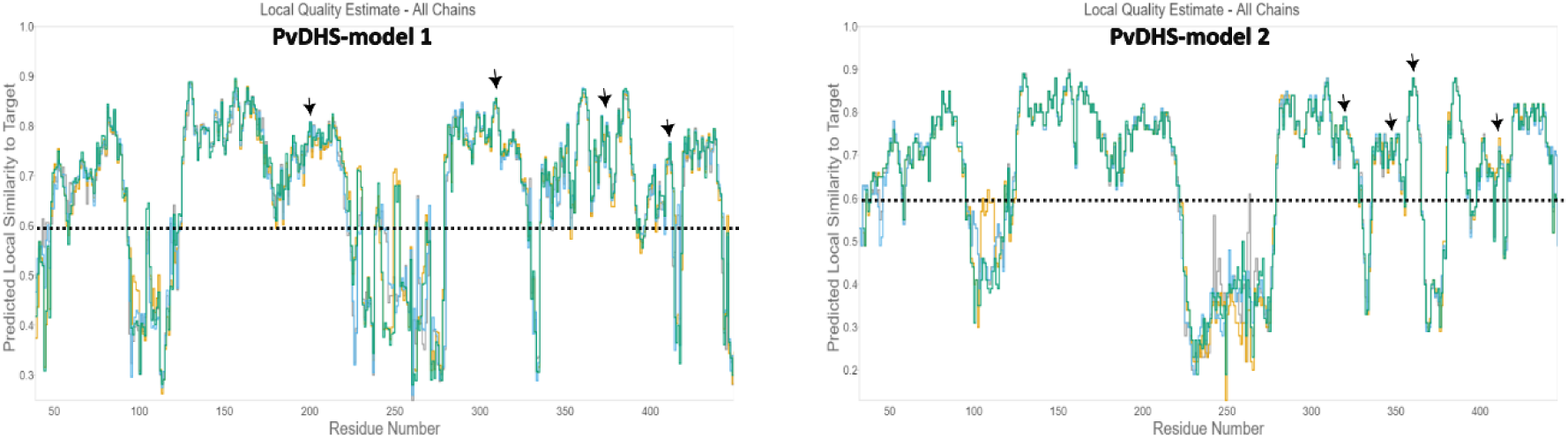
Local quality assessment of residues in the PvDHS-model 1 and PvDHS-model 2. The black dashed line indicates the threshold distinguishing poor- and high-quality local similarity regions. Arrows indicate the position of the residues within the binding sites.

**Figure S3.**
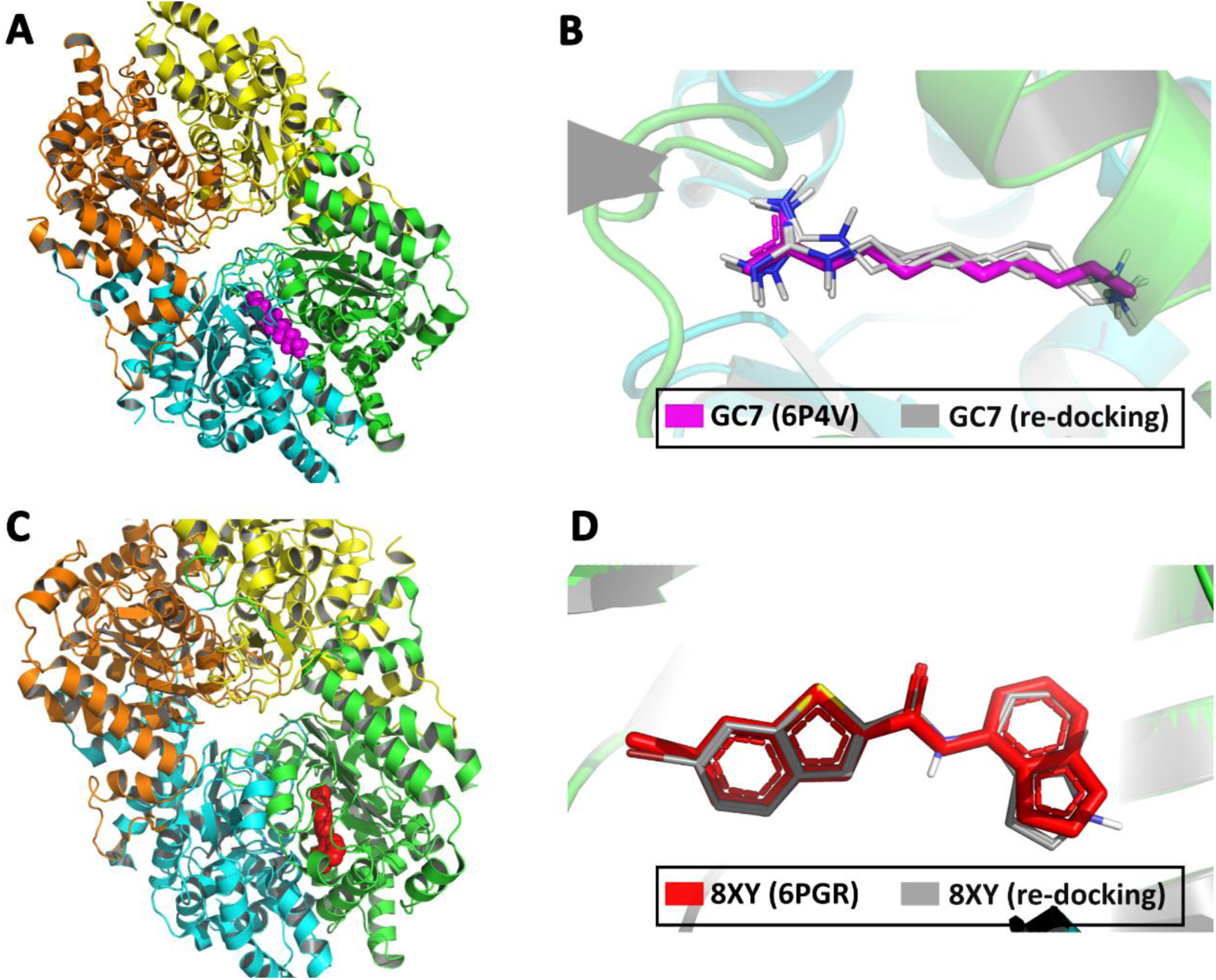
Alignment of the re-docked ligands (gray) with its original co-crystallized conformation. Zoomed-out view of GC7 (pink) and 8XY (red) disposition in the tetramer structures 6P4V (A) and PDB ID: 6PGR (C), respectively. B and D show the overlay of the original ligands, GC7 (pink) and 8XY (red) with the ligands after redocking (gray). Different colors represent individual chains in the structure.

**Figure S4.**
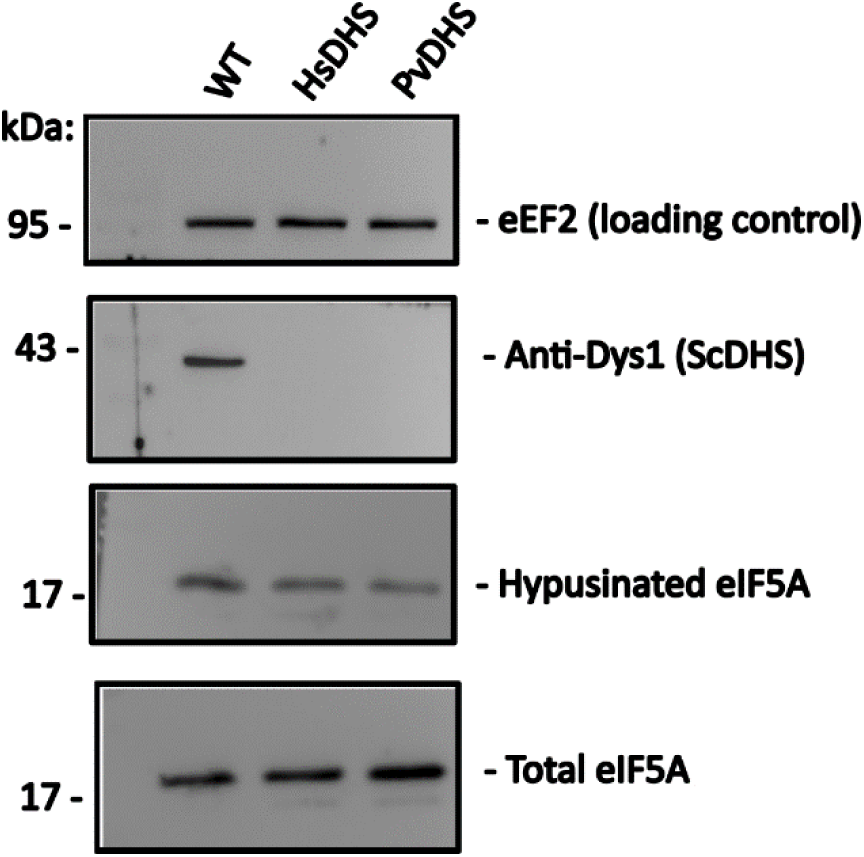
Hypusination of yeast eIF5A by *H. sapiens* and *P. vivax* DHS-complemented strains. Western blot detection of hypusinated eIF5A and total eIF5A from the *S. cerevisiae* strains wild-type, *dys1Δ::HsDHS* and *dys1Δ::PvDHS* (SFS01, SFS04 and SFS05, Table S2). An antibody specific for *S. cerevisiae* DHS (ScDHS) was used to show the deletion of the endogenous gene in the strains replaced by HsDHS and PvDHS.

**Figure S5.**
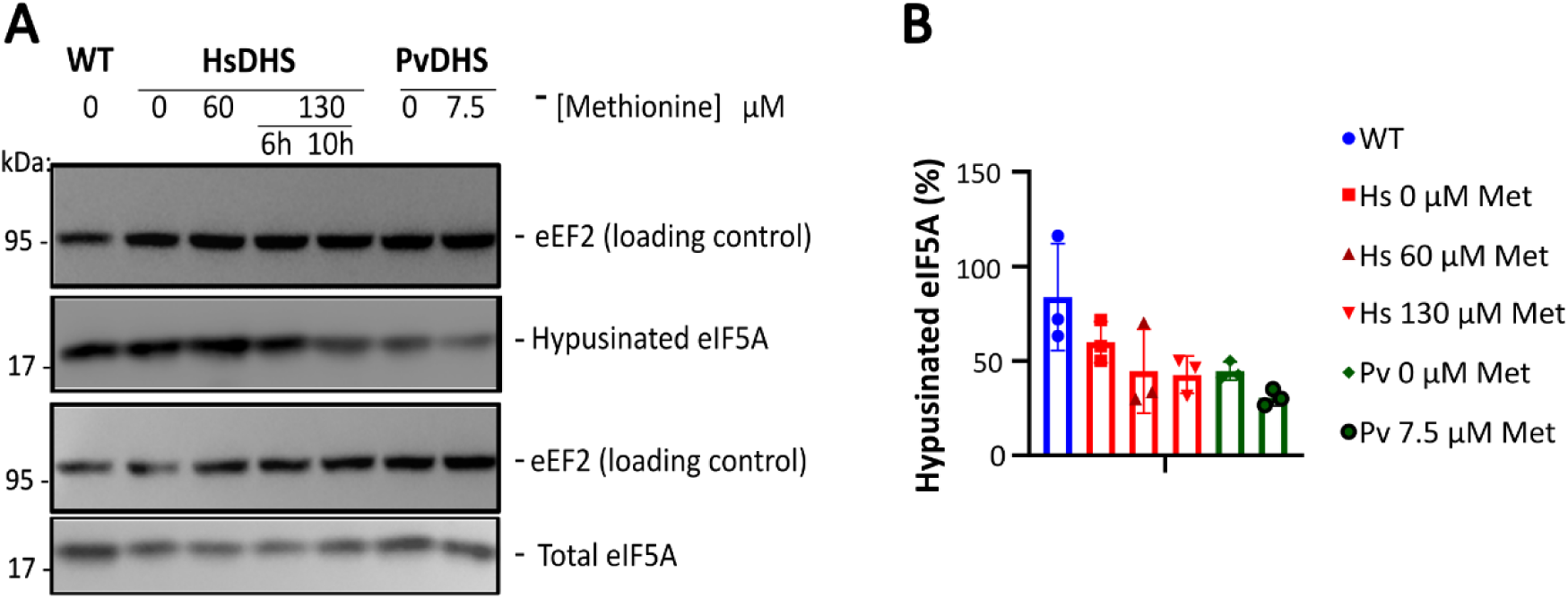
Hypusination of eIF5A under different DHS expression levels. (A) Western blot detection of hypusination levels after 10 h of methionine treatment (concentrations indicated in the figure). (B) Graph reporting the percentage of hypusinated eIF5A (hypusinated eIF5A/total eIF5A *100) under the conditions indicated in the figure. Bars represent mean ± standard deviation (n=3) of the percentage of hypusinated eIF5A.

**Figure S6.**
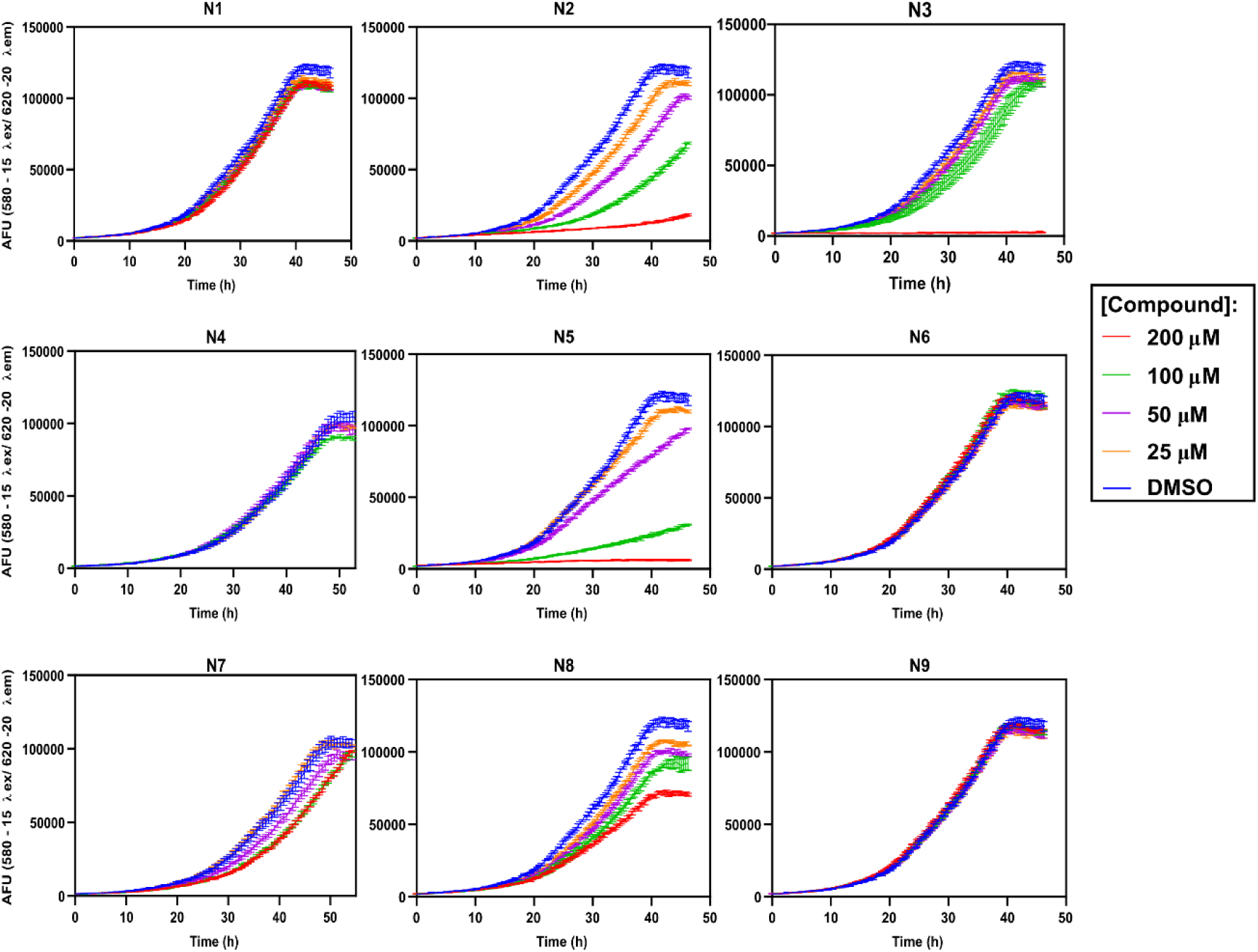
Growth of yeast DHS deleted strain complemented by PvDHS in the presence of compounds N1 to N9. The strain used was SFS05 (Table S2). The growth measurements were carried out in the Eve robot (see Materials and Methods) and it is given in arbitrary fluorescence units (AFU) (mean ± SD, n=4). Cell cultures were grown in SD–met and 1.25 % DMSO or 0, 25, 50, 100 and 200 μM concentrations of the respective compound tested (see the legend).

**Figure S7.**
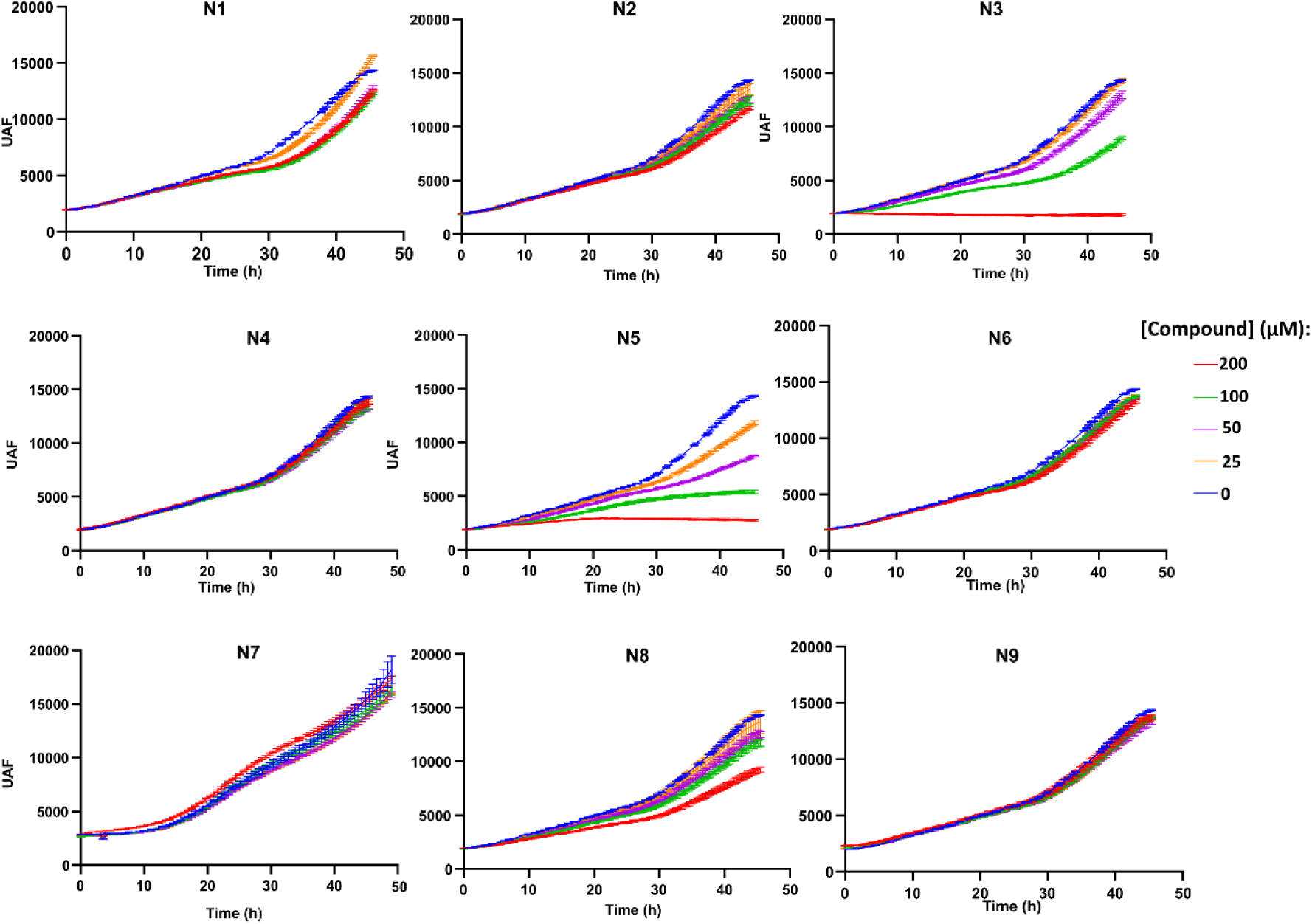
Growth of yeast DHS deleted strain complemented by HsDHS in the presence of compounds N1 to N9. The strain used was SFS04 (Table S2). The growth measurements, conducted in the Eve robot (see Materials and Methods) are presented in arbitrary fluorescence units (AFU) and shown as mean ± SD, n=4 replicates). Cell cultures were grown in SD with solvent alone (1.25 % DMSO) or varying concentrations (0, 25, 50, 100 and 200 μM) of the respective tested compound, as indicated in the legend.

**Figure S8.**
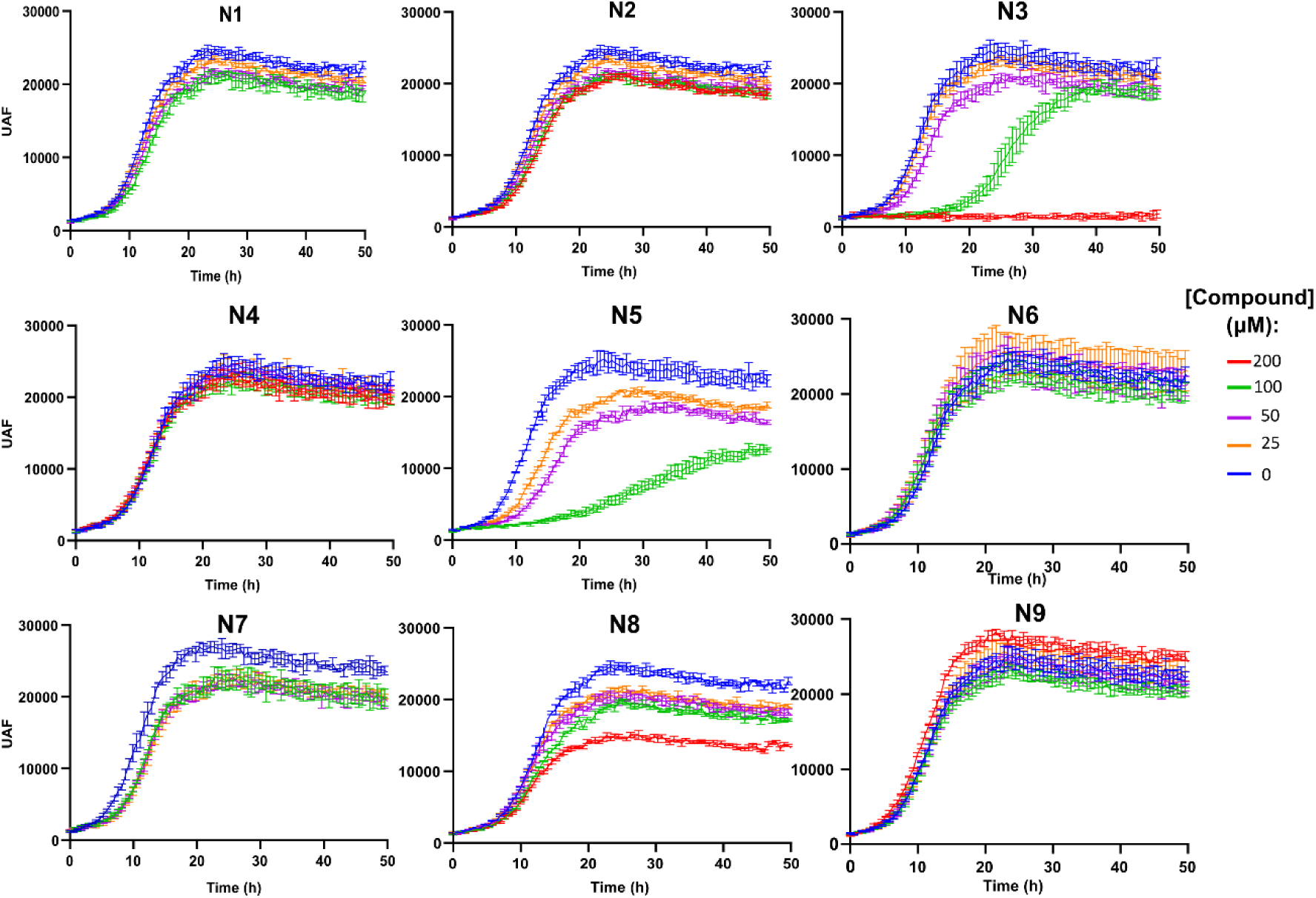
Growth of yeast wild type isogenic strain in the presence of compounds N1 to N9. The strain used was SFS01 (Table S2). The growth measurements were carried out in the Eve robot (see Materials and Methods) and it is given in arbitrary fluorescence units (AFU) (mean ± SD, n=4). Cell cultures were grown in SD and solvent alone (1.25 % DMSO) or varying concentrations or (25, 50, 100 and 200 μM) of the respective compound tested as indicated in the legend).

**Figure S9.**
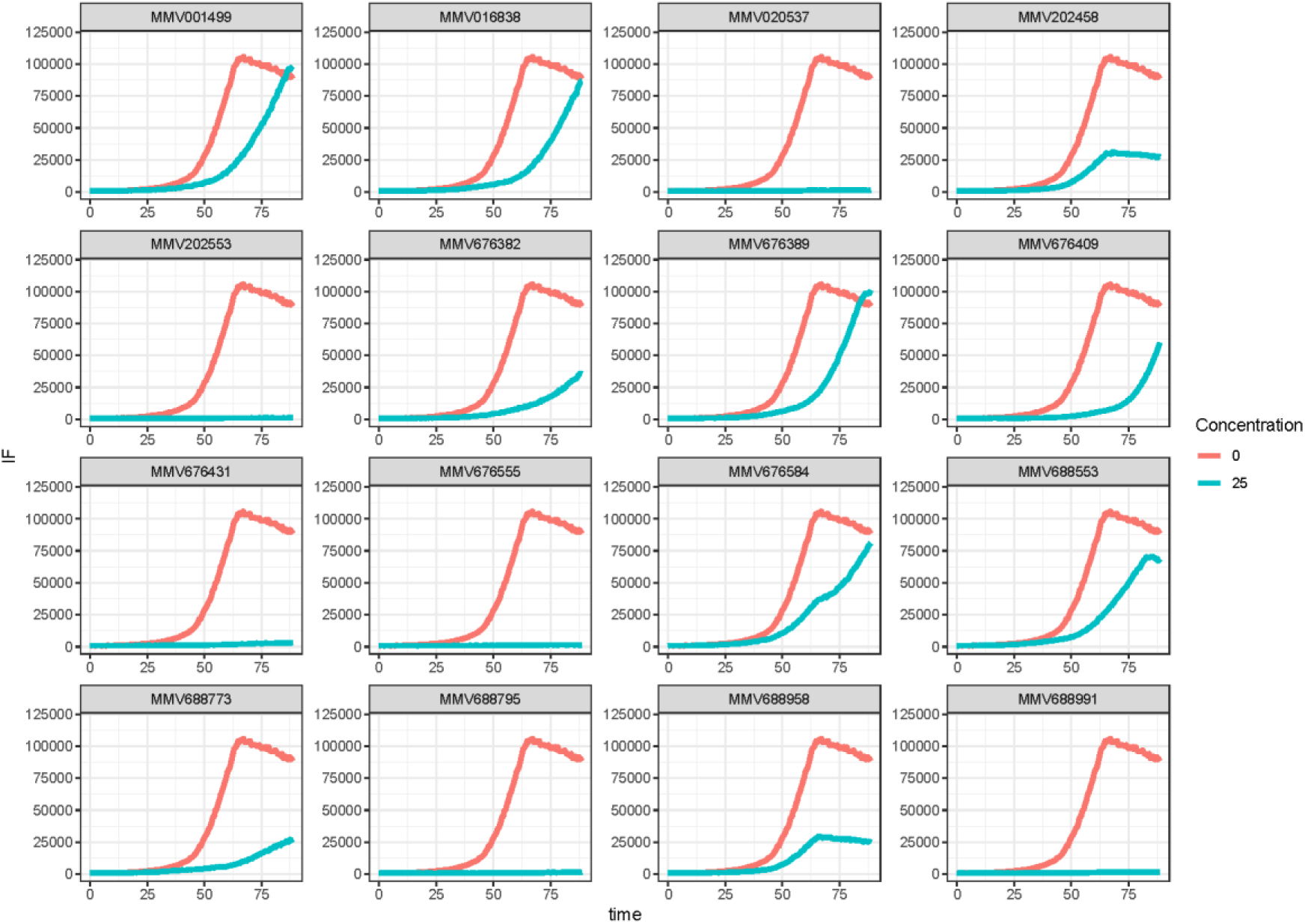
Growth reduction of PvDHS-complemented strain caused by compounds from the Pathogen Box. The strain used was SFS05 (Table S2). The growth measurements were carried out in the Eve robot (see Materials and Methods) and it is given in arbitrary fluorescence units (AFU) (mean ± SD, n=4). Cell cultures were grown in SD–met and 1.25 % DMSO or 25 μM of the respective compound tested (see the legend). The compounds are named according to the Pathogen box compound ID.

**Figure S10.**
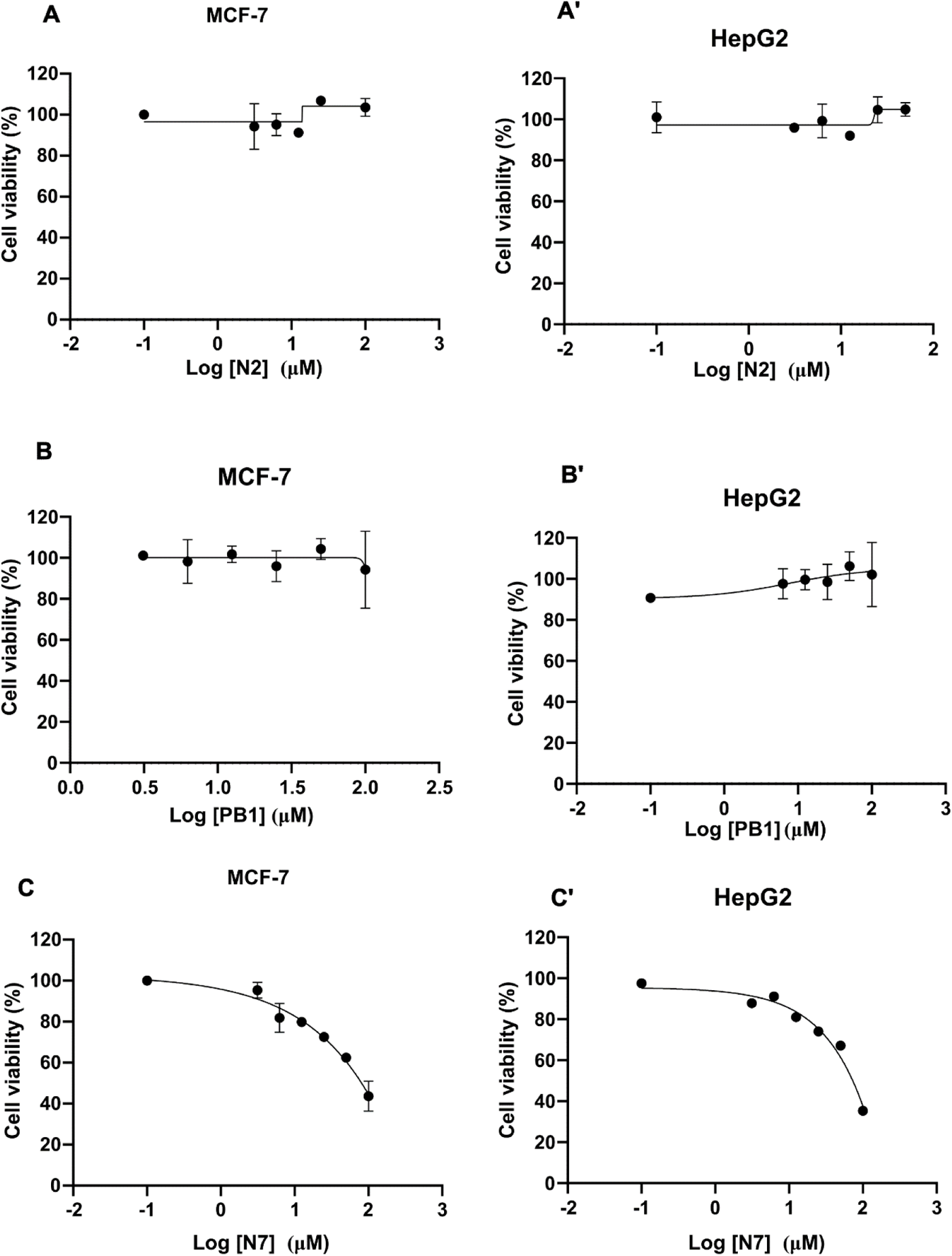
Cell viability analysis after treatment with compounds N2, N7 and PB1. The compounds were tested at concentrations ranging from 0.1 to 100 µM, for 24 h in MCF7 and HepG2 cells, as indicated in the panels). Cell viability (%) was calculated as the ratio between cell incubated with compound and cell incubated with DMSO. Each point represents the mean ± standard deviation, with n = 2 replicates.

## Supplementary Tables

**Table S1.** Plasmids used in this study.

| Plasmid | Features | Derived from | Source |
| --- | --- | --- | --- |
| pWS082 | sgRNA entry vector; *Amp<sup>R</sup>* | - | Tom Ellis (Addgene plasmid # 90516) |
| pWS082-CAN1 sgRNA | CAN1 sgRNA | pWS082 | This work |
| pWS082-dys1 sgRNA | DYS1 sgRNA | pWS082 | This work |
| pWS158 | Cas9 gap repair vector; *URA3*; *Kan<sup>R</sup>* | - | Tom Ellis (Addgene plasmid # 90517) |
| pWS172 | Cas9 gap repair vector; *HIS3*; *Kan<sup>R</sup>* | - | Tom Ellis (Addgene plasmid # 90519) |
| pCM188-MET3 | *CEN*; *URA3*; *Amp<sup>R</sup>*; *MET3pr* (methionine repressible) | - | [44] |
| pCM188-MET3-HsDHS | *Homo sapiens* DHS | pCM188-MET3 | This work |
| pCM188-MET3-PvDHS | *Plasmodium vivax* DHS | pCM188-MET3 | This work |
| yEp_CHERRY_HIS3 | *mCherry*; *HIS3* | - | [6] |
| yEp_Sapphire_HIS3 | *Sapphire*; *HIS3* | - | [6] |

**Table S2.**
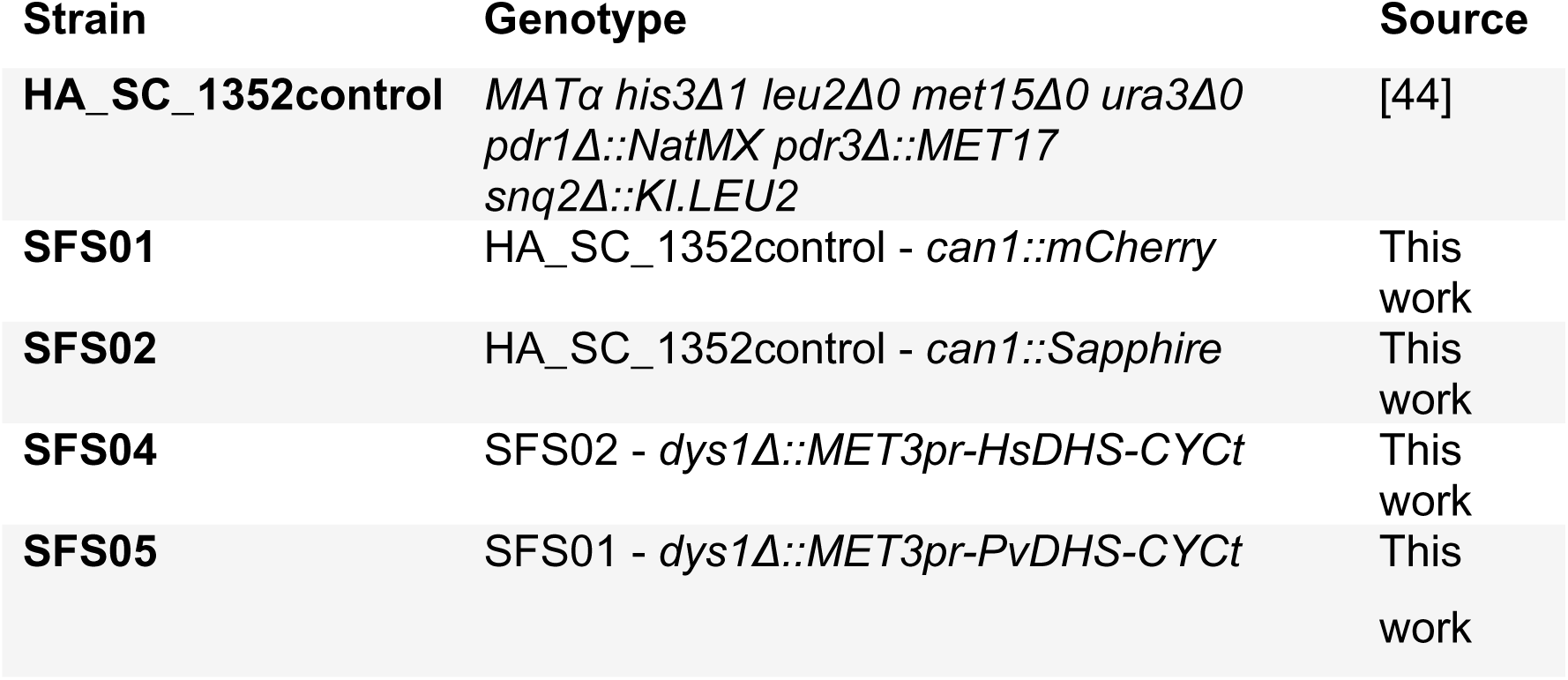
*Saccharomyces cerevisiae* strains used in this study.

**Table S3.**
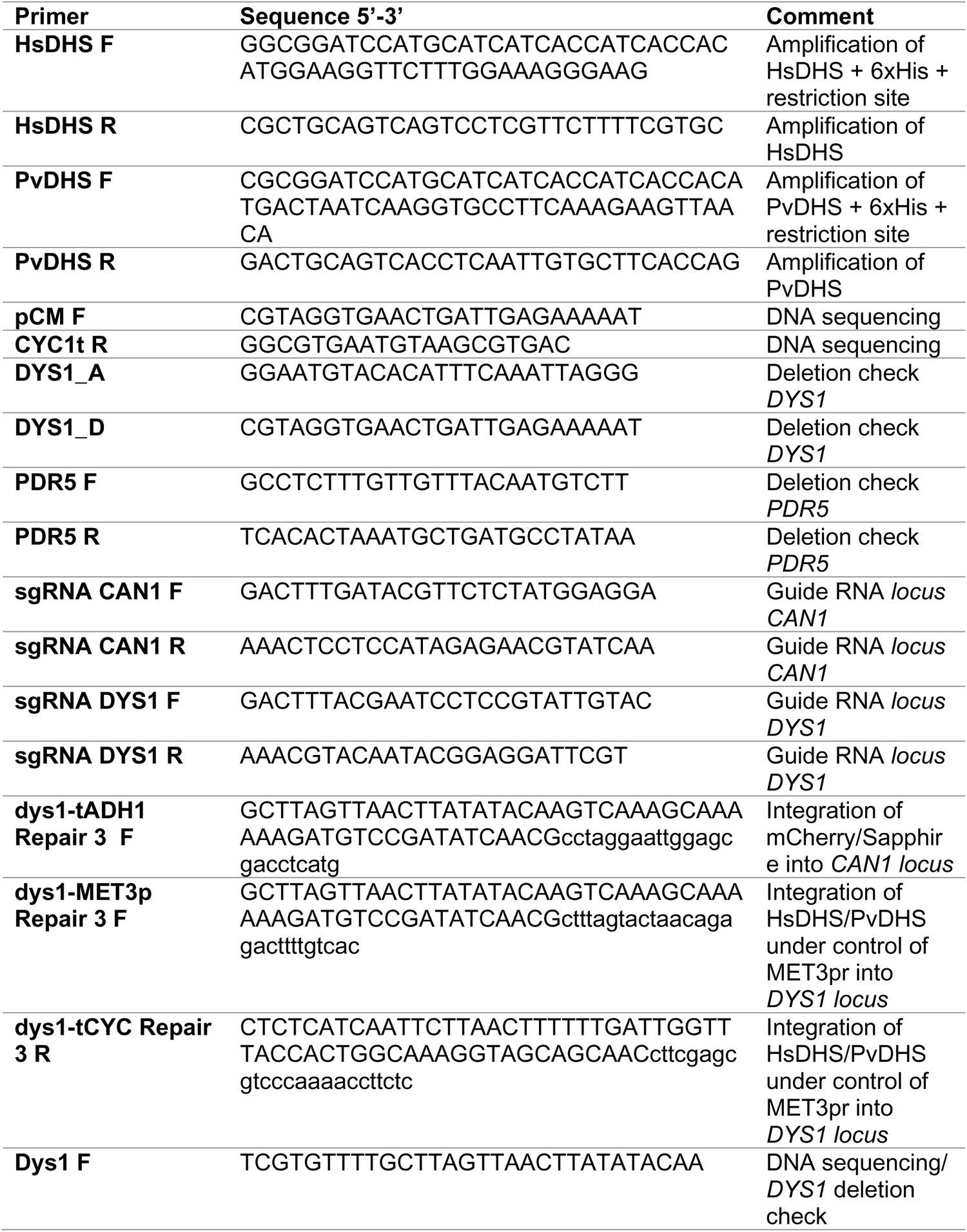
Oligonucleotides used in this study.

**Table S4.**
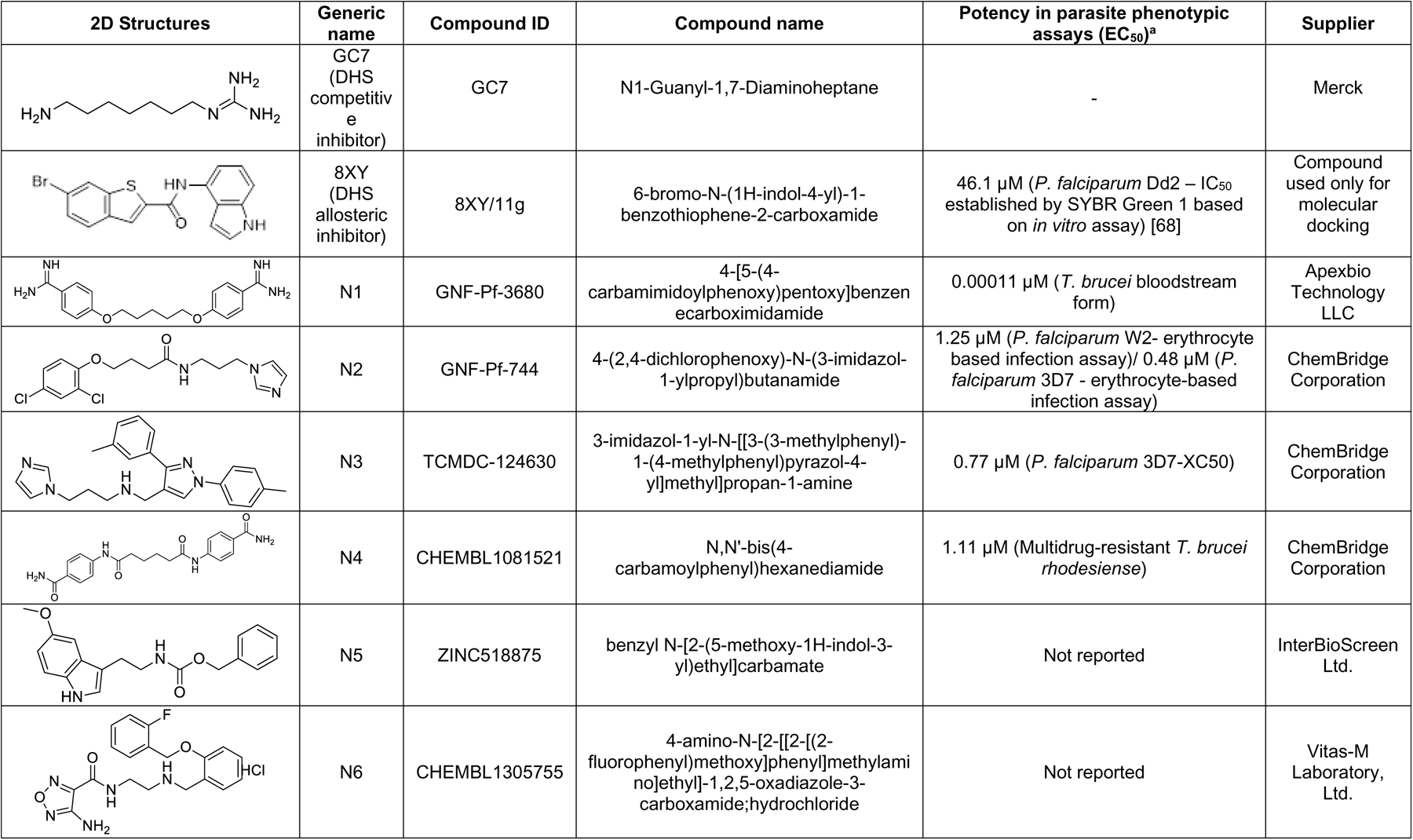
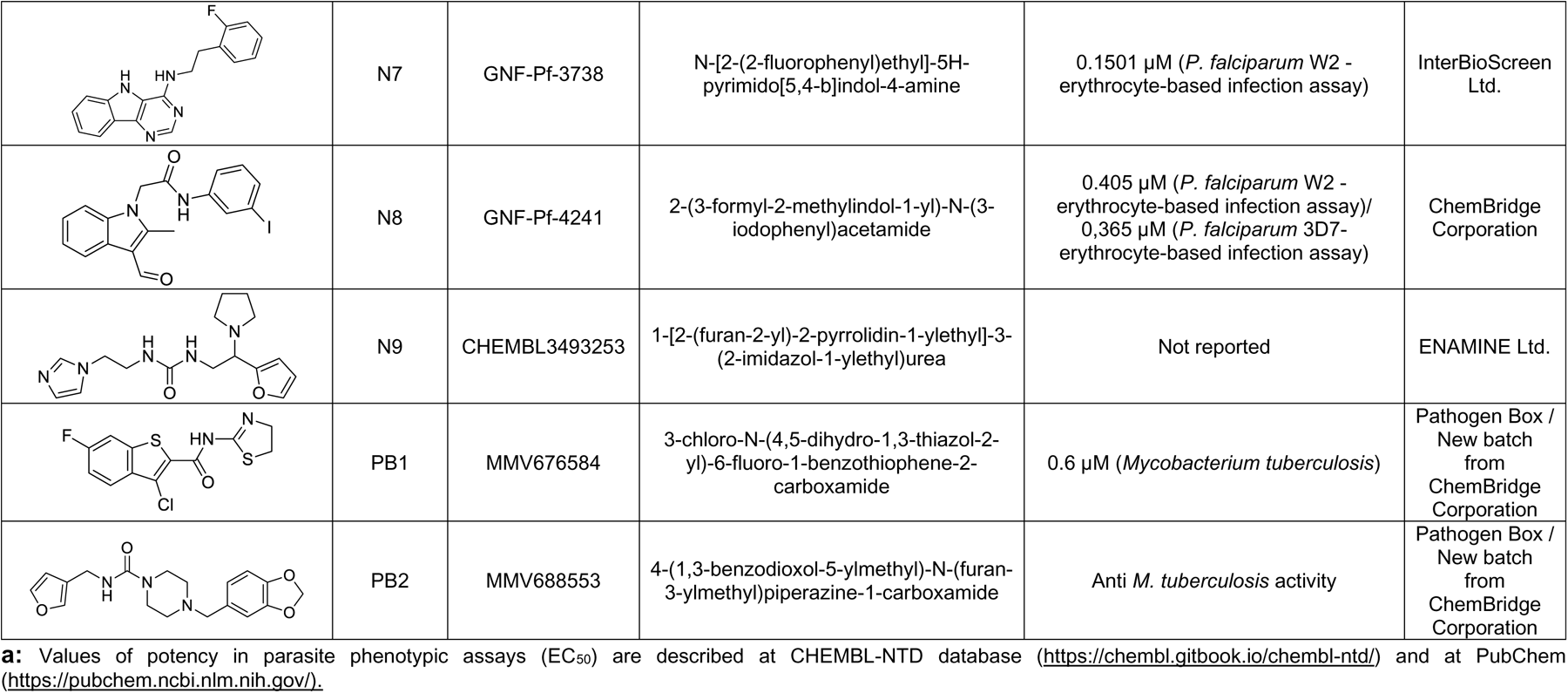
Synthetic compounds used in this study.

| 2D Structures | Generic name | Compound ID | Compound name | Potency in parasite phenotypic assays (EC<sub>50</sub>)<sup>a</sup> | Supplier |
| --- | --- | --- | --- | --- | --- |
|  | GC7<br>(DHS competitive inhibitor) | GC7 | N1-Guanyl-1,7-Diaminoheptane | - | Merck |
|  | 8XY<br>(DHS allosteric inhibitor) | 8XY/11g | 6-bromo-N-(1H-indol-4-yl)-1-benzothiophene-2-carboxamide | 46.1 μM (*P. falciparum* Dd2 – IC<sub>50</sub> established by SYBR Green 1 based on *in vitro* assay) [68] | Compound used only for molecular docking |
|  | N1 | GNF-Pf-3680 | 4-[5-(4-carbamimidoylphenoxy)pentoxy]benzenecarboximidamide | 0.00011 μM (*T. brucei* bloodstream form) | Apexbio Technology LLC |
|  | N2 | GNF-Pf-744 | 4-(2,4-dichlorophenoxy)-N-(3-imidazol-1-ylpropyl)butanamide | 1.25 μM (*P. falciparum* W2- erythrocyte based infection assay)/ 0.48 μM (*P. falciparum* 3D7 - erythrocyte-based infection assay) | ChemBridge Corporation |
|  | N3 | TCMDC-124630 | 3-imidazol-1-yl-N-[[3-(3-methylphenyl)-1-(4-methylphenyl)pyrazol-4-yl]methyl]propan-1-amine | 0.77 μM (*P. falciparum* 3D7-XC50) | ChemBridge Corporation |
|  | N4 | CHEMBL1081521 | N,N'-bis(4-carbamoylphenyl)hexanediamide | 1.11 μM (Multidrug-resistant *T. brucei rhodesiense*) | ChemBridge Corporation |
|  | N5 | ZINC518875 | benzyl N-[2-(5-methoxy-1H-indol-3-yl)ethyl]carbamate | Not reported | InterBioScreen Ltd. |
|  | N6 | CHEMBL1305755 | 4-amino-N-[2-[[2-[(2-fluorophenyl)methoxy]phenyl]methylamino]ethyl]-1,2,5-oxadiazole-3-carboxamide;hydrochloride | Not reported | Vitas-M Laboratory, Ltd. |
| | N7 | GNF-Pf-3738 | N-[2-(2-fluorophenyl)ethyl]-5H-pyrimido[5,4-b]indol-4-amine | 0.1501 μM (*P. falciparum* W2 - erythrocyte-based infection assay) | InterBioScreen Ltd. |
| | N8 | GNF-Pf-4241 | 2-(3-formyl-2-methylindol-1-yl)-N-(3-iodophenyl)acetamide | 0.405 μM (*P. falciparum* W2 - erythrocyte-based infection assay)/<br>0,365 μM (*P. falciparum* 3D7-erythrocyte-based infection assay) | ChemBridge Corporation |
|  | N9 | CHEMBL3493253 | 1-[2-(furan-2-yl)-2-pyrrolidin-1-ylethyl]-3-(2-imidazol-1-ylethyl)urea | Not reported | ENAMINE Ltd. |
| | PB1 | MMV676584 | 3-chloro-N-(4,5-dihydro-1,3-thiazol-2-yl)-6-fluoro-1-benzothiophene-2-carboxamide | 0.6 μM (*Mycobacterium tuberculosis*) | Pathogen Box /<br>New batch<br>from<br>ChemBridge<br>Corporation |
|  | PB2 | MMV688553 | 4-(1,3-benzodioxol-5-ylmethyl)-N-(furan-3-ylmethyl)piperazine-1-carboxamide | Anti *M. tuberculosis* activity | Pathogen Box /<br>New batch<br>from<br>ChemBridge<br>Corporation |

**a:** Values of potency in parasite phenotypic assays (EC<sub>50</sub>) are described at ChEMBL-NTD database (https://chembl.gitbook.io/chembl-ntd/) and at PubChem (https://pubchem.ncbi.nlm.nih.gov/).

---

**Text S1. Synthetic DHS genes with codon usage optimized for expression in *Saccharomyces cerevisiae***.

The coding region from each gene was flanked by *BamH*I and *Pst*I restriction sites (highlighted in yellow).

**HsDHS**

GGATCCATGGAAGGTTCTTTGGAAAGGGAAGCTCCAGCTGGTGCATTGGC TGCTGTTTTGAAACATTCTTCTACTTTGCCACCAGAATCTACCCAAGTTAGA GGTTACGATTTTAACAGAGGTGTTAACTACAGAGCTTTGTTGGAAGCTTTT GGTACTACTGGTTTCCAAGCTACTAATTTCGGTAGAGCTGTTCAACAAGTT AACGCCATGATTGAAAAGAAGTTGGAACCATTGTCTCAAGACGAAGATCAA CATGCTGATTTGACTCAATCTAGAAGGCCATTGACATCTTGCACTATTTTCT TGGGTTACACCTCCAACTTGATCTCTTCTGGTATTAGAGAAACCATCAGAT ACTTGGTCCAACACAACATGGTTGATGTTTTGGTTACTACAGCTGGTGGTG TTGAAGAGGATTTGATTAAGTGTTTGGCTCCAACTTACTTGGGTGAATTTTC CTTGAGAGGCAAAGAATTGAGAGAGAACGGTATTAACAGAATCGGCAATTT GTTGGTCCCAAACGAAAATTACTGCAAGTTCGAAGATTGGTTGATGCCAAT CTTGGATCAAATGGTCATGGAACAAAACACCGAAGGTGTTAAGTGGACTCC ATCTAAAATGATTGCCAGATTGGGCAAAGAAATCAACAATCCAGAATCCGT TTATTACTGGGCCCAAAAGAATCATATCCCAGTTTTTTCACCAGCTTTGACC GATGGTTCTTTAGGTGATATGATCTTCTTCCACTCCTACAAAAATCCAGGTT TGGTTTTGGATATCGTCGAGGACTTGAGATTGATTAACACCCAAGCTATTTT CGCTAAGTGCACCGGTATGATTATCTTAGGTGGTGGTGTAGTCAAACATCA TATTGCTAATGCTAACTTGATGAGAAACGGTGCTGATTACGCTGTTTACATT AACACTGCTCAAGAATTCGACGGTTCTGATTCAGGTGCTAGACCAGATGAA GCTGTTTCTTGGGGTAAAATTAGAGTTGATGCTCAACCAGTTAAGGTTTAC GCTGATGCTTCTTTGGTTTTCCCATTATTGGTTGCTGAAACCTTCGCTCAAA AGATGGATGCTTTTATGCACGAAAAGAACGAGGACTGATGACTGCAGGGT ACCTGGAG

**PvDHS**

GGATCCATGACTAATCAAGGTGCCTTCAAAGAAGTTAACAAGATCAGGTCT GAATCCGATGATGGTGAATCTTCTGACGAAAAGTCTGGTATTGAAGATGCC AAGTCATCCGTTTTCGTTAAGTCCAACAAAATCCCAGAAAACACCGATGTT GTTAAGGGTATCAACTTCGAAGAAGAAGTCAACTTGCACCAATTCGTTAAC CAGTATAAGTACATGGGTTTCCAAGCTACCAACTTAGGTATTGGTATCGAT GAGGTCAACAAGATGATCCATTTTAAGTATGCTGAAGGTGGTGAAGGTACT CAAGATGGTCATGATAATGATCACGATCAAGATTCCGATGACGAAAGACAA GCTTTGCCAAAGAAAAAGAAGTGCTTGATCTGGTTGTCTTTCACCTCCAAT ATGATCTCTTCTGGTTTGAGAGAAATCTTCGTCTACCTGGCTAAGAAAAAG TTCATCGATGTTGTCGTTACTACTGCTGGTGGTGTTGAAGAGGATATTATC AAGTGTTTCTCCAAGACTTACTTGGGCGATTTTAACTTGAACGGTAAGAAG TTGAGAAAGAAAGGTTGGAACAGAATCGGCAATTTGATTGTCCCAAACGAT AACTACTGCAAGTTCGAAGATTGGTTGCAGCCTTTGTTGAACAAGATGTTG CATGAACAGAACCGTAAGAACGAAGAGTTGTTCTTGAGAAAGTTGGACAAA AGACGTAGAGGTGGTGGTCATGGTGGTGAAAGGGAACCACCATCTCCACC TCCACATACACCACATGCTCCATCACCACCAAGTCCATGTGATTCTTCAGA TGAAGATGAATCCGACATGTTCTACTTGTCTCCATCTGAATTCATCGACAAATTGGGCGAAGAAATCAACGACGAATCTTCTTTGATATACTGGTGCCACAAG AACGATATTCCAGTTTTTTGTCCAGGTTTGACCGATGGTTCTTTGGGTGATA ATTTGTTCTTCCACAATTACGGCAAGAAGATCAAAAACAACCTGATCCTGG ATATCGTCAAGGACATCAAGAAGATTAACTCATTGGCTCTGAACTGCAAGA AGTCCGGTATTATCATTTTAGGTGGTGGCTTGCCAAAACATCATGTCTGTA ATGCTAACTTGATGAGAAACGGTGCTGATTTCGCTGTTTACGTTAATACTG CTAACGAATACGACGGTTCTGATTCTGGTGCTAACACTACTGAAGCTTTGT CTTGGGGTAAAATCAAAGCTGGTCATACCAACAACCACGTTAAGGTTTTTG GTGATGCTACCATTTTGTTCCCATTGATGGTTTTGAACACCTTCTACTTGCA TGACAGAGGTGGTAGACATAATTCTGGTGAAGCACAATTGAGGTGACTGG AG

## DATA AVAILABILITY

The PvDHS-docked best poses, a table with description of the compounds selected by virtual screening and a table reporting the datasets tested *in silico* are available as free download at Zenodo.org, https://doi.org/10.5281/zenodo.10006188.

DNA sequencing files from cloning constructs and genes integrated into the *CAN1* locus, mCherry, Sapphire, and into the *DYS1* locus, HsDHS and PvDHS are available at Zenodo.org, https://doi.org/10.5281/zenodo.10053251.

## References

1. 1. WHO. World malaria report 2022 2022. Available from: https://www.who.int/teams/global-malaria-programme/reports/world-malaria-report-2022.

2. Carlson CJ, Bannon E, Mendenhall E, Newfield T, Bansal S. Rapid range shifts in African *Anopheles* mosquitoes over the last century. Biol Lett. 2023;19:2022036520220365. doi: 10.1098/rsbl.2022.0365.

3. Fairlamb AH, Gow NA, Matthews KR, Waters AP. Drug resistance in eukaryotic microorganisms. Nat Microbiol. 2016;1(7):16092. Epub 20160624. doi: 10.1038/nmicrobiol.2016.92. PubMed PMID: 27572976; PubMed Central PMCID: PMCPMC5215055.

4. Battle KE, Baird JK. The global burden of *Plasmodium vivax* malaria is obscure and insidious. PLoS Med. 2021;18(10):e1003799. Epub 20211007. doi: 10.1371/journal.pmed.1003799. PubMed PMID: 34618814; PubMed Central PMCID: PMCPMC8496786.

5. Bilsland E, Bean DM, Devaney E, Oliver SG. Yeast-based high-throughput screens to identify novel compounds active against *Brugia malay*i. PLoS Negl Trop Dis. 2016;10(1):e0004401. doi: 10.1371/journal.pntd.0004401. PubMed PMID: 26812604; PubMed Central PMCID: PMC4727890.

6. Bilsland E, Sparkes A, Williams K, Moss HJ, de Clare M, Pir P, et al. Yeast-based automated high-throughput screens to identify anti-parasitic lead compounds. Open Biol. 2013;3(2):120158. doi: 10.1098/rsob.120158. PubMed PMID: 23446112.

7. Silva SF, Klippel AH, Ramos PZ, Santiago ADS, Valentini SR, Bengtson MH, et al. Structural features and development of an assay platform of the parasite target deoxyhypusine synthase of *Brugia malayi* and *Leishmania major*. PLoS Negl Trop Dis. 2020;14(10):e0008762. Epub 20201012. doi: 10.1371/journal.pntd.0008762. PubMed PMID: 33044977; PubMed Central PMCID: PMCPMC7581365.

8. Frearson JA, Wyatt PG, Gilbert IH, Fairlamb AH. Target assessment for antiparasitic drug discovery. Trends Parasitol. 2007;23(12):589–95. Epub 20071024. doi: 10.1016/j.pt.2007.08.019. PubMed PMID: 17962072; PubMed Central PMCID: PMCPMC2979298.

9. Wator E, Wilk P, Biela A, Rawski M, Zak KM, Steinchen W, et al. Cryo-EM structure of human eIF5A-DHS complex reveals the molecular basis of hypusination-associated neurodegenerative disorders. Nat Commun. 2023;14(1):1698. Epub 20230327. doi: 10.1038/s41467-023-37305-2. PubMed PMID: 36973244; PubMed Central PMCID: PMCPMC10042821.

10. Park MH, Nishimura K, Zanelli CF, Valentini SR. Functional significance of eIF5A and its hypusine modification in eukaryotes. Amino Acids. 2010;38(2):491–500. Epub 20091208. doi: 10.1007/s00726-009-0408-7. PubMed PMID: 19997760; PubMed Central PMCID: PMCPMC2829442.

11. Park MH, Joe YA, Kang KR. Deoxyhypusine synthase activity is essential for cell viability in the yeast *Saccharomyces cerevisiae*. J Biol Chem. 1998;273(3):1677–83. doi: 10.1074/jbc.273.3.1677. PubMed PMID: 9430712.

12. Nishimura K, Lee SB, Park JH, Park MH. Essential role of eIF5A-1 and deoxyhypusine synthase in mouse embryonic development. Amino Acids. 2012;42(2-3):703–10. Epub 20110818. doi: 10.1007/s00726-011-0986-z. PubMed PMID: 21850436; PubMed Central PMCID: PMCPMC3220921.

13. Wator E, Wilk P, Grudnik P. Half way to hypusine-structural basis for substrate recognition by human deoxyhypusine synthase. Biomolecules. 2020;10(4). Epub 20200330. doi: 10.3390/biom10040522. PubMed PMID: 32235505; PubMed Central PMCID: PMCPMC7226451.

14. Aroonsri A, Posayapisit N, Kongsee J, Siripan O, Vitsupakorn D, Utaida S, et al. Validation of *Plasmodium falciparum* deoxyhypusine synthase as an antimalarial target. PeerJ. 2019;7:e6713. Epub 20190417. doi: 10.7717/peerj.6713. PubMed PMID: 31024761; PubMed Central PMCID: PMCPMC6475138.

15. Nguyen S, Jones DC, Wyllie S, Fairlamb AH, Phillips MA. Allosteric activation of trypanosomatid deoxyhypusine synthase by a catalytically dead paralog. J Biol Chem. 2013;288(21):15256–67. doi: 10.1074/jbc.M113.461137. PubMed PMID: 23525104; PubMed Central PMCID: PMC3663545.

16. Quintas-Granados LI, Carvajal Gamez BI, Villalpando JL, Ortega-Lopez J, Arroyo R, Azuara-Liceaga E, et al. Bifunctional activity of deoxyhypusine synthase/hydroxylase from *Trichomonas vaginalis*. Biochimie. 2016;123:37–51. Epub 20150926. doi: 10.1016/j.biochi.2015.09.027. PubMed PMID: 26410361.

17. Coni S, Serrao SM, Yurtsever ZN, Di Magno L, Bordone R, Bertani C, et al. Blockade of EIF5A hypusination limits colorectal cancer growth by inhibiting MYC elongation. Cell Death Dis. 2020;11(12):1045. Epub 20201210. doi: 10.1038/s41419-020-03174-6. PubMed PMID: 33303756; PubMed Central PMCID: PMCPMC7729396.

18. He LR, Zhao HY, Li BK, Liu YH, Liu MZ, Guan XY, et al. Overexpression of eIF5A-2 is an adverse prognostic marker of survival in stage I non-small cell lung cancer patients. Int J Cancer. 2011;129(1):143–50. doi: 10.1002/ijc.25669. PubMed PMID: 20830705.

19. Li L, Li X, Zhang Q, Ye T, Zou S, Yan J. EIF5A expression and its role as a potential diagnostic biomarker in hepatocellular carcinoma. J Cancer. 2021;12(16):4774–9. Epub 20210611. doi: 10.7150/jca.58168. PubMed PMID: 34234848; PubMed Central PMCID: PMCPMC8247388.

20. Sfakianos AP, Raven RM, Willis AE. The pleiotropic roles of eIF5A in cellular life and its therapeutic potential in cancer. Biochem Soc Transact. 2022;50(6):1885–95. doi: 10.1042/BST20221035. PubMed PMID: 36511302; PubMed Central PMCID: PMCPMC9788402.

21. Hardbower DM, Asim M, Luis PB, Singh K, Barry DP, Yang C, et al. Ornithine decarboxylase regulates M1 macrophage activation and mucosal inflammation via histone modifications. Proc Natl Acad Sci USA. 2017;114(5):E751-E60. Epub 20170117. doi: 10.1073/pnas.1614958114. PubMed PMID: 28096401; PubMed Central PMCID: PMCPMC5293075.

22. Mastracci TL, Colvin SC, Padgett LR, Mirmira RG. Hypusinated eIF5A is expressed in the pancreas and spleen of individuals with type 1 and type 2 diabetes. PLoS One. 2020;15(3):e0230627. Epub 20200324. doi: 10.1371/journal.pone.0230627. PubMed PMID: 32208453; PubMed Central PMCID: PMCPMC7092972.

23. Maier B, Tersey SA, Mirmira RG. Hypusine: a new target for therapeutic intervention in diabetic inflammation. Discov Med. 2010;10(50):18–23. PubMed PMID: 20670594.

24. Padgett LR, Shinkle MR, Rosario S, Stewart TM, Foley JR, Casero RA, Jr., et al. Deoxyhypusine synthase mutations alter the post-translational modification of eukaryotic initiation factor 5A resulting in impaired human and mouse neural homeostasis. HGG Adv. 2023;4(3):100206. Epub 20230518. doi: 10.1016/j.xhgg.2023.100206. PubMed PMID: 37333770; PubMed Central PMCID: PMCPMC10275725.

25. Liang Y, Piao C, Beuschel CB, Toppe D, Kollipara L, Bogdanow B, et al. eIF5A hypusination, boosted by dietary spermidine, protects from premature brain aging and mitochondrial dysfunction. Cell Rep. 2021;35(2):108941. doi: 10.1016/j.celrep.2021.108941. PubMed PMID: 33852845.

26. Liu J, Henao-Mejia J, Liu H, Zhao Y, He JJ. Translational regulation of HIV-1 replication by HIV-1 Rev cellular cofactors Sam68, eIF5A, hRIP, and DDX3. J Neuroimmune Pharmacol. 2011;6(2):308–21. Epub 20110301. doi: 10.1007/s11481-011-9265-8. PubMed PMID: 21360055.

27. Jakus J, Wolff EC, Park MH, Folk JE. Features of the spermidine-binding site of deoxyhypusine synthase as derived from inhibition studies. Effective inhibition by bis- and mono-guanylated diamines and polyamines. J Biol Chem. 1993;268(18):13151–9. PubMed PMID: 8514754.

28. Tanaka Y, Kurasawa O, Yokota A, Klein MG, Ono K, Saito B, et al. Discovery of novel allosteric inhibitors of deoxyhypusine synthase. J Med Chem. 2020;63(6):3215–26. Epub 20200316. doi: 10.1021/acs.jmedchem.9b01979. PubMed PMID: 32142284.

29. Liu KL, Li XY, Wang DP, Xue WH, Qian XH, Li YH, et al. Novel allosteric inhibitors of deoxyhypusine synthase against malignant melanoma: design, synthesis, and biological evaluation. J Med Chem. 2021;64(18):13356–72. Epub 20210902. doi: 10.1021/acs.jmedchem.1c00582. PubMed PMID: 34473510.

30. Zhou QY, Tu CY, Shao CX, Wang WK, Zhu JD, Cai Y, et al. GC7 blocks epithelial-mesenchymal transition and reverses hypoxia-induced chemotherapy resistance in hepatocellular carcinoma cells. Am J Transl Res. 2017;9(5):2608–17. Epub 20170515. PubMed PMID: 28560008; PubMed Central PMCID: PMCPMC5446540.

31. Altamura F, Rajesh R, Catta-Preta CMC, Moretti NS, Cestari I. The current drug discovery landscape for trypanosomiasis and leishmaniasis: Challenges and strategies to identify drug targets. Drug Dev Res. 2022;83(2):225–52. Epub 20200406. doi: 10.1002/ddr.21664. PubMed PMID: 32249457.

32. Denny PW. Yeast: bridging the gap between phenotypic and biochemical assays for high-throughput screening. Expert Opin Drug Discov. 2018;13(12):1153–60. Epub 20181016. doi: 10.1080/17460441.2018.1534826. PubMed PMID: 30326751.

33. Denny PW, Steel PG. Yeast as a potential vehicle for neglected tropical disease drug discovery. J Biomol Screen. 2015;20(1):56–63. Epub 20140813. doi: 10.1177/1087057114546552. PubMed PMID: 25121554.

34. Cao Y, Sun C, Wen H, Wang M, Zhu P, Zhong M, et al. A yeast-based drug discovery platform to identify *Plasmodium falciparum* type II NADH dehydrogenase inhibitors. Antimicrob Agents Chemother. 2021. doi: 10.1128/AAC.02470-20. PubMed PMID: 33722883.

35. Norcliffe JL, Mina JG, Alvarez E, Cantizani J, de Dios-Anton F, Colmenarejo G, et al. Identifying inhibitors of the *Leishmania* inositol phosphorylceramide synthase with antiprotozoal activity using a yeast-based assay and ultra-high throughput screening platform. Sci Rep. 2018;8(1):3938. Epub 20180302. doi: 10.1038/s41598-018-22063-9. PubMed PMID: 29500420; PubMed Central PMCID: PMCPMC5834442.

36. Land H, Humble MS. YASARA: A tool to obtain structural guidance in biocatalytic investigations. Methods Mol Biol. 2018;1685:43–67. doi: 10.1007/978-1-4939-7366-8_4. PubMed PMID: 29086303.

37. Umland TC, Wolff EC, Park MH, Davies DR. A new crystal structure of deoxyhypusine synthase reveals the configuration of the active enzyme and of an enzyme.NAD.inhibitor ternary complex. J Biol Chem. 2004;279(27):28697–705. Epub 20040420. doi: 10.1074/jbc.M404095200. PubMed PMID: 15100216.

38. Afanador GA, Tomchick DR, Phillips MA. Trypanosomatid deoxyhypusine synthase activity is dependent on shared active-Sste complementation between pseudoenzyme paralogs. Structure. 2018;26(11):1499–512 e5. Epub 20180906. doi: 10.1016/j.str.2018.07.012. PubMed PMID: 30197036; PubMed Central PMCID: PMCPMC6221947.

39. Pandey R, Kumar R, Gupta P, Mohmmed A, Tewari R, Malhotra P, et al. High throughput *in silico* identification and characterization of *Plasmodium falciparum* PRL phosphatase inhibitors. J Biomol Struct Dyn. 2018;36(13):3531-Epub 20171106. doi: 10.1080/07391102.2017.1392365. PubMed PMID: 29039247.

40. Solis EJ, Pandey JP, Zheng X, Jin DX, Gupta PB, Airoldi EM, et al. Defining the essential function of yeast Hsf1 reveals a compact transcriptional program for maintaining eukaryotic proteostasis. Mol Cell. 2018;69(3):534. doi: 10.1016/j.molcel.2018.01.021. PubMed PMID: 29395071; PubMed Central PMCID: PMCPMC5813686.

41. Shi XP, Yin KC, Ahern J, Davis LJ, Stern AM, Waxman L. Effects of N1-guanyl-1,7-diaminoheptane, an inhibitor of deoxyhypusine synthase, on the growth of tumorigenic cell lines in culture. Biochim Biophys Acta. 1996;1310(1):119–26. doi: 10.1016/0167-4889(95)00165-4. PubMed PMID: 9244184.

42. Bilsland E, Pir P, Gutteridge A, Johns A, King RD, Oliver SG. Functional expression of parasite drug targets and their human orthologs in yeast. PLoS Negl Trop Dis. 2011;5(10):e1320. doi: 10.1371/journal.pntd.0001320. PubMed PMID: 21991399; PubMed Central PMCID: PMC3186757.

43. Da Silva NA, Srikrishnan S. Introduction and expression of genes for metabolic engineering applications in *Saccharomyces cerevisiae*. FEMS Yeast Res. 2012;12(2):197–214. Epub 20120112. doi: 10.1111/j.1567-1364.2011.00769.x. PubMed PMID: 22129153.

44. Alalam H, Sigurdardóttir S, Bourgard C, Tiukova IA, King RD, Grøtli M, et al. A genetic trap in yeast for inhibitors of the SARS-CoV-2 main protease. mSystems. 2021;6:e01087–21. doi: 10.1128/mSystems.01087-21

45. Buechel ER, Pinkett HW. Transcription factors and ABC transporters: from pleiotropic drug resistance to cellular signaling in yeast. FEBS Lett. 2020;594(23):3943–64. Epub 20201107. doi: 10.1002/1873-3468.13964. PubMed PMID: 33089887.

46. Piotrowski JS, Li SC, Deshpande R, Simpkins SW, Nelson J, Yashiroda Y, et al. Functional annotation of chemical libraries across diverse biological processes. Nat Chem Biol. 2017;13(9):982–93. doi: 10.1038/nchembio.2436. PubMed PMID: 28759014.

47. Ryan OW, Skerker JM, Maurer MJ, Li X, Tsai JC, Poddar S, et al. Selection of chromosomal DNA libraries using a multiplex CRISPR system. eLife. 2014;3. Epub 20140819. doi: 10.7554/eLife.03703. PubMed PMID: 25139909; PubMed Central PMCID: PMCPMC4161972.

48. Mao X, Hu Y, Liang C, Lu C. *MET3* promoter: a tightly regulated promoter and its application in construction of conditional lethal strain. Curr Microbiol. 2002;45(1):37–40. doi: 10.1007/s00284-001-0046-0. PubMed PMID: 12029525.

49. 49. Deposited Set 2: Novartis-GNF Malaria Box Dataset (hits from P. falciparum whole-cell screening). Available from: https://chembl.gitbook.io/chembl-ntd/#deposited-set-2-20th-may-2010-novartis-gnf-malaria-box-dataset-hits-from-p.-falciparum-whole-cell-sc.

50. Mugumbate G, Mendes V, Blaszczyk M, Sabbah M, Papadatos G, Lelievre J, et al. Target identification of *Mycobacterium tuberculosis* phenotypic hits using a concerted chemogenomic, biophysical, and structural approach. Front Pharmacol. 2017;8:681. Epub 20170926. doi: 10.3389/fphar.2017.00681. PubMed PMID: 29018348; PubMed Central PMCID: PMCPMC5623190.

51. 51. Burke D, Dawson D, Stearns T. Methods in yeast genetics: a Cold Spring Harbor Laboratory course manual.: Cold Spring Harbor Laboratory Press; 2000.

52. Ausubel FM, editor. Current Protocols in Molecular Biology. New York: John Wiley & Sons; 2020.

53. Park MH, Wolff EC, Lee YB, Folk JE. Antiproliferative effects of inhibitors of deoxyhypusine synthase. Inhibition of growth of Chinese hamster ovary cells by guanyl diamines. J Biol Chem. 1994;269(45):27827–32. PubMed PMID: 7961711.

54. Benkert P, Biasini M, Schwede T. Toward the estimation of the absolute quality of individual protein structure models. Bioinformatics. 2011;27(3):343–50. Epub 20101205. doi: 10.1093/bioinformatics/btq662. PubMed PMID: 21134891; PubMed Central PMCID: PMCPMC3031035.

55. Sastry GM, Adzhigirey M, Day T, Annabhimoju R, Sherman W. Protein and ligand preparation: parameters, protocols, and influence on virtual screening enrichments. J Comput Aided Mol Des. 2013;27(3):221–34. Epub 20130412. doi: 10.1007/s10822-013-9644-8. PubMed PMID: 23579614.

56. Jacobson MP, Friesner RA, Xiang Z, Honig B. On the role of the crystal environment in determining protein side-chain conformations. J Mol Biol. 2002;320(3):597–608. doi: 10.1016/S0022-2836(02)00470-9. PubMed PMID: 12096912.

57. Jacobson MP, Pincus DL, Rapp CS, Day TJ, Honig B, Shaw DE, et al. A hierarchical approach to all-atom protein loop prediction. Proteins. 2004;55(2):351–67. doi: 10.1002/prot.10613. PubMed PMID: 15048827.

58. Lu C, Wu C, Ghoreishi D, Chen W, Wang L, Damm W, et al. OPLS4: Improving force field accuracy on challenging regimes of chemical space. J Chem Theory Comput. 2021;17(7):4291–300. Epub 20210607. doi: 10.1021/acs.jctc.1c00302. PubMed PMID: 34096718.

59. Shelley JC, Cholleti A, Frye LL, Greenwood JR, Timlin MR, Uchimaya M. Epik: a software program for pK(a) prediction and protonation state generation for drug-like molecules. J Comput Aided Mol Des. 2007;21(12):681–91. Epub 20070927. doi: 10.1007/s10822-007-9133-z. PubMed PMID: 17899391.

60. Friesner RA, Murphy RB, Repasky MP, Frye LL, Greenwood JR, Halgren TA, et al. Extra precision glide: docking and scoring incorporating a model of hydrophobic enclosure for protein-ligand complexes. J Med Chem. 2006;49(21):6177–96. doi: 10.1021/jm051256o. PubMed PMID: 17034125.

61. Wichapong K, Rohe A, Platzer C, Slynko I, Erdmann F, Schmidt M, et al. Application of docking and QM/MM-GBSA rescoring to screen for novel Myt1 kinase inhibitors. J Chem Inf Model. 2014;54(3):881–93. Epub 20140213. doi: 10.1021/ci4007326. PubMed PMID: 24490903.

62. DiCarlo JE, Norville JE, Mali P, Rios X, Aach J, Church GM. Genome engineering in *Saccharomyces cerevisiae* using CRISPR-Cas systems. Nucl Acids Res. 2013;41(7):4336–43. Epub 20130304. doi: 10.1093/nar/gkt135. PubMed PMID: 23460208; PubMed Central PMCID: PMCPMC3627607.

63. Marillonnet S, Grutzner R. Synthetic DNA assembly using Golden Gate cloning and the hierarchical modular cloning pipeline. Curr Protoc Mol Biol. 2020;130(1):e115. doi: 10.1002/cpmb.115. PubMed PMID: 32159931.

64. Gietz RD, Schiestl RH. Large-scale high-efficiency yeast transformation using the LiAc/SS carrier DNA/PEG method. Nat Protocol. 2007;2(1):38–41. doi: 10.1038/nprot.2007.15. PubMed PMID: 17401336.

65. Akhmetov A, Laurent JM, Gollihar J, Gardner EC, Garge RK, Ellington AD, et al. Single-step precision genome editing in yeast using CRISPR-Cas9. Bio Protoc. 2018;8(6). doi: 10.21769/BioProtoc.2765. PubMed PMID: 29770349; PubMed Central PMCID: PMCPMC5951413.

66. Sprouffske K, Wagner A. Growthcurver: an R package for obtaining interpretable metrics from microbial growth curves. BMC Bioinformatics. 2016;17:172. Epub 20160419. doi: 10.1186/s12859-016-1016-7. PubMed PMID: 27094401; PubMed Central PMCID: PMCPMC4837600.

67. 67. Lazar I. Gelanalyzer. Available from: http://gelanalyzer.com/?i=1.

68. Aroonsri A, Wongsombat C, Shaw P, Franke S, Przyborski J, Kaiser A. Investigation of an allosteric deoxyhypusine synthase inhibitor in *P. falciparum*. Molecules. 2022;27(8). Epub 20220411. doi: 10.3390/molecules27082463. PubMed PMID: 35458660; PubMed Central PMCID: PMCPMC9030622.

